# Integrator governs NF-κB dwell time on chromatin to drive inflammatory transcription

**DOI:** 10.64898/2026.09.25.754463

**Authors:** Pradeep Kumar Reddy Cingaram, Jingyin Yue, Fan Liu, Helena Gomes Dos Santos, Gabriel S. Gaidosh, Simone Sidoli, Benjamin A Garcia, Fan Lai, Mingjiang Xu, Omar Lopez, Guilherme Miura Lavezzo, Felipe Beckedorff, Stephen D. Nimer, Ramin Shiekhattar

## Abstract

NF-κB must remain bound to chromatin long enough to activate innate immune genes, but what stabilizes this transcriptionally productive state is unknown. Here, we identify the Integrator complex as an essential determinant of NF-κB chromatin residence in mouse and human cells. Integrator associated with NF-κB and co-occupied its genomic targets. Single-molecule tracking revealed that Integrator markedly prolonged the chromatin residence of RELA molecules in a low-mobility state. Catalytically inactive INTS11-E203Q fully restored NF-κB–dependent transcription, separating this chromatin-retention function from RNA cleavage. Conditional *Ints11* deletion abolished TLR4-dependent dendritic-cell activation in adult mice and impaired LPS-induced transcription in mouse embryonic fibroblasts. Loss of Integrator in human cancer cells similarly suppressed TNFα-induced inflammatory gene expression. These findings uncover a noncatalytic function for Integrator that stabilizes NF-κB at enhancers, complementing its established catalytic control of RNA polymerase II pause release. By coupling signal recognition at enhancers to productive elongation, Integrator provides a molecular solution to a central challenge of multicellularity: converting transient extracellular cues into coherent gene-expression programs across complex chromatin landscapes.

## Introduction

Nuclear factor kappa B (NF-κB) comprises a family of five transcription factors (TFs) that form various homo or heterodimers, which bind to consensus DNA sequences at regulatory regions of genes responsive to inflammatory stimuli(*1*). Activation of canonical NF-κB, composed of a RELA/p50 heterodimer, is rapidly and transiently induced by various stimuli, including pro-inflammatory cytokines (TNF-α, IL1b), bacterial and viral products (lipopolysaccharides (LPS), double-stranded RNAs, among many others(*2*). The primary step for NF-κB activation entails the release of NF-κB, from the inhibitory IκB proteins. Stimulus-induced activation of inhibitors of IκB kinase (IKK) leads to phosphorylation of the inhibitor IκB-, which binds and retains inactive NF-κB in the cytoplasm(*3*). Upon its phosphorylation, IκB-is targeted for K48-linked ubiquitylation and subsequent proteasomal degradation(*4*). The resulting free NF-κB can translocate to the nucleus where it binds to cognate recognition sites at distal or promoter-proximal enhancers of inflammation promoting genes(*5–7*).

While most NF-κB binding sites are found in nucleosome free regions (NFRs) or open chromatin, others are obscured by nucleosomes and require additional regulatory mechanisms to provide chromatin accessibility(*8, 9*). Recent studies have pointed to a class of lineage specifying transcription factors (TF) termed, pioneer factors, that have the ability to interact with their DNA binding sites imbedded in nucleosomes. Among the first examples of pioneer factors, FOXA and GATA were shown to cooperatively associate with nucleosomal sites(*10, 11*). It has been hypothesized that binding of pioneer factors to chromatin allows for DNA interactions of other TFs unable to find their DNA cognate sites on nucleosomes(*11, 12*). Interestingly, while NF-κB predominantly binds open chromatin, detailed analyses of NF-κB binding *in vivo* indicates that a stable association of NF-κB with DNA requires additional cofactors providing increased binding energy(*13–19*).

In line with this observation, we find that the multi-protein complex, Integrator, co-occupies NF-κB binding sites and is essential for NF-κB chromatin association and responsiveness to a variety of inflammatory stimuli. Single molecule tracking experiments revealed that Integrator is critical for enhanced residence time of the bound state of NF-κB to its cognate sites. We show that this function of Integrator is independent of it is catalytic endonuclease activity. Therefore, Integrator fulfill two key roles in transcriptional regulation. Through its catalytic activity, Integrator plays a critical role in regulating RNAPII pause release to promote transcriptional elongation(*20–22*). While in this study, we broaden the scope of Integrator function beyond its catalytic activity to the regulation of innate immune response by licensing NF-κB chromatin residence.

## Results

### Integrator orchestrates transcriptional responsiveness to LPS and TNF-α

Since Integrator plays an essential role in immediate early gene responsiveness (*20, 21, 23, 24*), we assessed Integrator role in NF-κB transcriptional induction of inflammatory genes. Mouse embryonic fibroblasts (MEFs) were previously used to assess NF-κB transcriptional dynamics(*25*). We isolated MEFs from wild type mice or mice with homozygous deletion of *Ints11* (Fig. S1A). MEFs were then treated with 200 ng/ml LPS for 1 hour following which changes in gene expression were assessed using RNA sequencing (Fig. 1A and B). LPS stimulation elicited a robust activation of inflammatory genes in MEFs which was highly dampened following deletion of INTS11 (Fig. 1A-D, Fig. S1B, Table 1). We validated loss of LPS responsiveness in three LPS responsive genes using real time polymerase chain reaction (PCR) following INTS11 deletion (Fig. 1E-G). These results highlight a critical role for INTS11 in LPS-induced activation of IEGs in MEFs.

**Figure 1.**
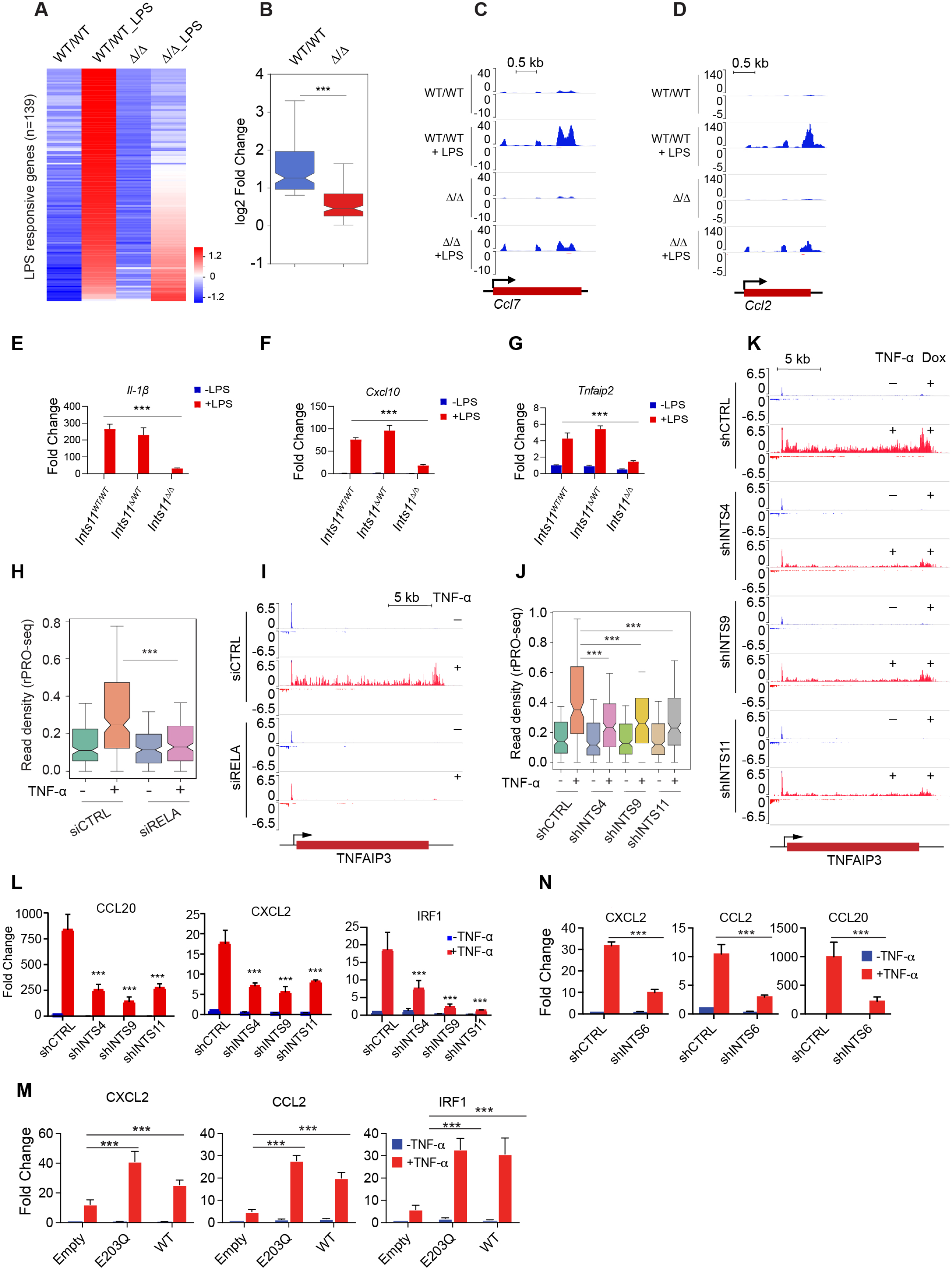
Integrator regulates transcriptional responses to LPS and TNF-α. **(A)** RNA-seq heat map showing 139 LPS-responsive genes in *Ints11* ^WT/WT^ and *Ints11* ^Δ/Δ^ mouse embryonic fibroblasts (MEFs), before and after LPS stimulation. **(B)** Box plot showing the distribution of log2 fold changes in expression of the 139 LPS-responsive genes in *Ints11* ^WT/WT^ and *Ints11* ^Δ/Δ^ MEFs. ***P < 0.0001, two-tailed Student’s *t* test. **(C and D)** Representative RNA-seq genome-browser tracks at the *Ccl7* **(C)** and *Ccl2* **(D)** loci in *Ints11* ^WT/WT^ and *Ints11* ^Δ/Δ^ MEFs before and after LPS stimulation. **(E to G)** RT-qPCR analysis of the LPS-responsive genes *Il1b* **(E)**, *Cxcl10* **(F)**, and *Tnfaip2* **(G)** in *Ints11* ^WT/WT^, *Ints11* ^Δ/WT^, and *Ints11* ^Δ/Δ^ MEFs before and after LPS stimulation. ***P < 0.001. **(H)** Box plots showing PRO-seq read density within the gene bodies of TNF-α responsive genes in HeLa cells treated with control siRNA (siCTRL) or RELA-targeting siRNA (siRELA), before and after TNF-α stimulation. ***P < 0.001. **(I)** Representative rPRO-seq genome-browser tracks at the *TNFAIP3* locus in siCTRL and siRELA HeLa cells before and after TNF-α stimulation. **(J)** Box plots showing rPRO-seq read density within the gene bodies of TNF-α responsive genes in HeLa shCTRL, shINTS4, shINTS9, and shINTS11 cells before and after TNF-α stimulation. ***P < 0.001. **(K)** Representative rPRO-seq genome-browser tracks at the *TNFAIP3* locus in Dox-treated HeLa shCTRL, shINTS4, shINTS9, and shINTS11 cells before and after TNF-α stimulation. **(L)** RT-qPCR analysis of the TNF-α responsive genes *CCL20*, *CXCL2*, and *IRF1* in HeLa shCTRL, shINTS4, shINTS9, and shINTS11 cells before and after TNF-α stimulation. Statistical comparisons were performed between TNF-α stimulated shCTRL cells and the corresponding Integrator-depleted cells using an unpaired two-tailed Student’s *t* test. ***P < 0.001. **(M)** RT-qPCR analysis of the TNF-α responsive genes *CXCL2*, *CCL2*, and *IRF1* in Dox-treated INTS11-depleted cells reconstituted with empty vector, WT-INTS11, or the E203Q-INTS11 catalytic mutant, before and after TNF-α stimulation. Statistical comparisons of TNF-α stimulated cells expressing empty vector with those expressing E203Q-INTS11 or WT-INTS11 are indicated. ***P < 0.001, unpaired two-tailed Student’s *t* test. **(N)** RT-qPCR analysis of *CXCL2*, *CCL2*, and *CCL20* expression in shCTRL and shINTS6 HeLa cells before and after TNF-α stimulation. Statistical comparisons were performed between TNF-α stimulated shCTRL and shINTS6 cells using an unpaired two-tailed Student’s *t* test. ***P < 0.001.

**Figure S1.**
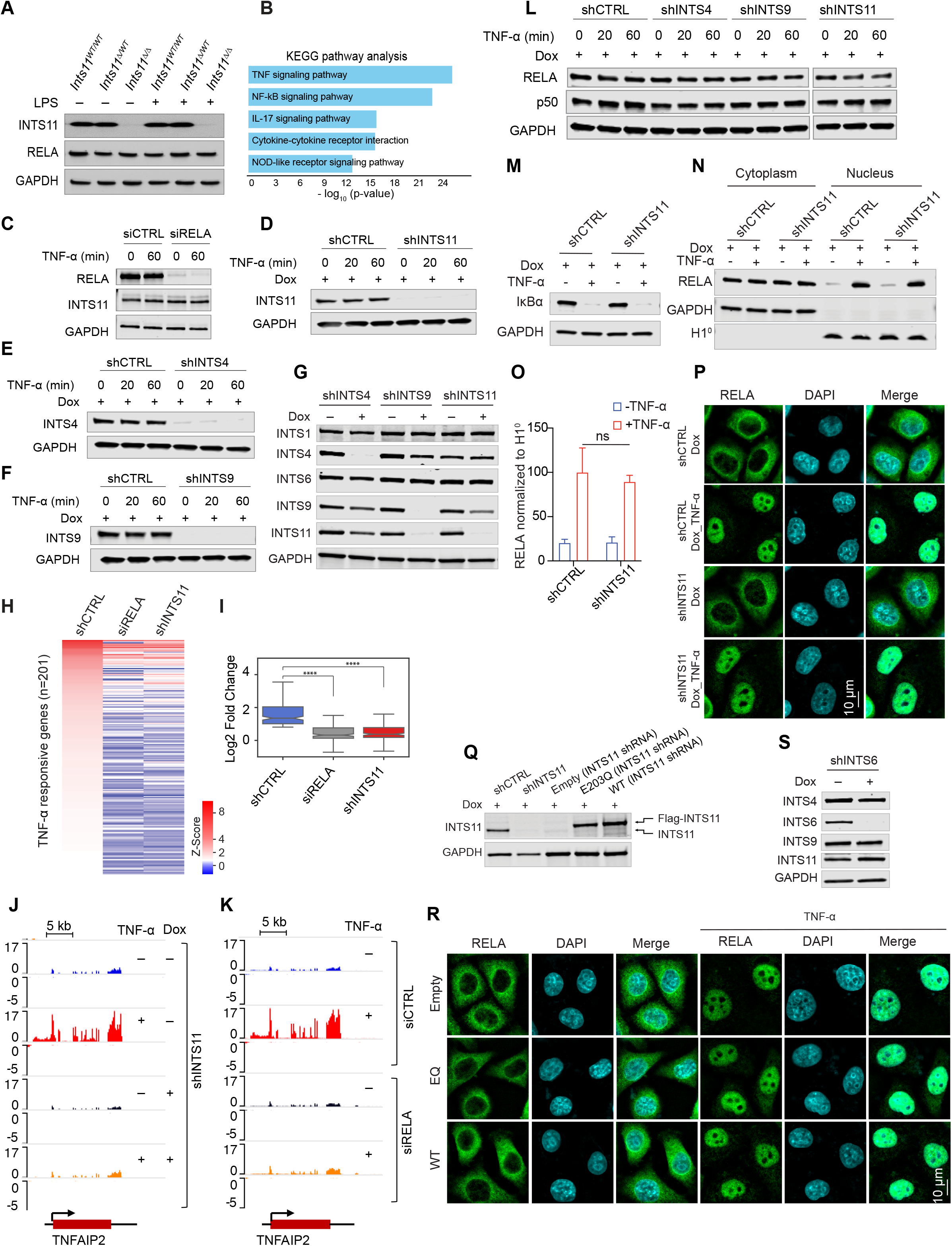
Depletion of Integrator impairs activation of NF-κB dependent genes. **(A)** Immunoblot analysis of INTS11 and RELA in *Ints11* WT/WT, *Ints11* Δ/WT, and *Ints11* Δ/Δ MEFs before and after LPS stimulation. GAPDH served as a loading control. **(B)** KEGG pathway enrichment analysis of genes up-regulated in MEFs following LPS stimulation. Enrichment significance is shown as −log10(*P* value). **(C)** Immunoblot validation of RELA depletion in siCTRL and siRELA HeLa cells before and 60 min after TNF-α stimulation. INTS11 abundance is also shown; GAPDH served as a loading control. **(D to F)** Immunoblot validation of Dox-induced depletion of INTS11 **(D)**, INTS4 **(E)**, and INTS9 **(F)** in the corresponding HeLa shRNA cell lines during a TNF-α stimulation time course. GAPDH served as a loading control. **(G)** Immunoblot analysis of Integrator subunits INTS1, INTS4, INTS6, INTS9, and INTS11 in shINTS4, shINTS9, and shINTS11 HeLa cells cultured in the absence or presence of Dox. GAPDH served as a loading control. **(H)** Heat map showing the transcriptional responses of 201 TNF-α responsive genes in HeLa shCTRL, siRELA, and shINTS11 cells following 60 min of TNF-α stimulation. Genes are ordered according to their TNF-α responsiveness, and values are displayed as Z scores. **(I)** Box plots showing the distribution of log2 fold changes for the 201 TNF-α responsive genes in shCTRL, siRELA, and shINTS11 cells. ****P < 0.0001, Student’s *t* test. **(J)** Representative RNA-seq genome-browser tracks at the *TNFAIP2* locus in shINTS11 HeLa cells under the indicated Dox and TNF-α treatment conditions. **(K)** Representative RNA-seq genome-browser tracks at the *TNFAIP2* locus in siRELA HeLa cells before and after TNF-α stimulation. **(L)** Immunoblot analysis of RELA and p50 in Dox-treated shCTRL, shINTS4, shINTS9, and shINTS11 HeLa cells at 0, 20, and 60 min after TNF-α stimulation. GAPDH served as a loading control. **(M)** Immunoblot analysis of IκBα in Dox-treated shCTRL and shINTS11 HeLa cells before and after TNF-α stimulation. GAPDH served as a loading control. **(N)** Immunoblot analysis of RELA in cytoplasmic and nuclear fractions of Dox-treated shCTRL and shINTS11 HeLa cells before and after TNF-α stimulation. GAPDH and histone H1.0 served as cytoplasmic and nuclear fraction markers, respectively. **(O)** Quantification of nuclear RELA abundance normalized to histone H1.0 in shCTRL and shINTS11 cells before and after TNF-α stimulation. Data are presented as mean ± SD; ns, not significant. **(P)** Representative immunofluorescence images showing RELA (green) and DAPI (cyan) in Dox-treated shCTRL and shINTS11 cells before and after TNF-α stimulation. Scale bar, 10 μm. **(Q)** Immunoblot analysis confirming INTS11 depletion and re-expression of shRNA-resistant FLAG-tagged WT-INTS11 or E203Q-INTS11. Cells expressing shCTRL, shINTS11, empty vector, E203Q-INTS11, or WT-INTS11 are shown as indicated. Endogenous and FLAG-tagged INTS11 are indicated; GAPDH served as a loading control. **(R)** Representative immunofluorescence images showing RELA (green) and DAPI (cyan) under the indicated control, knockdown, and INTS11 reconstitution conditions, before and after TNF-α stimulation. Scale bar, 10 μm. **(S)** Immunoblot analysis of INTS4, INTS6, INTS9, and INTS11 in shINTS6 cells cultured in the absence or presence of Dox. GAPDH served as a loading control.

We next dissected the molecular underpinning of Integrator function in NF-κB signaling in HeLa cells, a cervical cancer cell line, that allows for large-scale molecular analyses including nascent transcriptome measurements and displays a robust responsiveness to TNF-α stimulation. Importantly, NF-κB signaling plays an important role in human cervical cancer(*26*). We deplete the canonical NF-κB subunit RELA or Integrator subunits, INTS11, INTS9 or INTS4 (Fig. S1C-G) and assess changes in nascent gene expression using rapid precision nuclear run-on sequencing (rPRO-seq). Loss of RELA or Integrator subunits significantly diminish the expression of inflammatory genes following TNF-α stimulation (Fig. 1H-K). These include genes such as CCL20, CXCL2 and IRF1 (Fig. 1L, and Table 2). Further analyses of inflammatory gene expression by RNA-seq or real time-PCR confirm the loss of TNF-α responsiveness (Fig. 1L, Fig. S1H-K, Table 2). Importantly, depletion of INTS11 does not affect the TNF-α induced translocation of RELA into the nucleus or the degradation of IκB-, confirming that acute depletion of INTS11 does not disrupt the cytoplasmic to nuclear translocation of NF-κB (Fig. S1M-P). These results establish a critical and a specific role for Integrator in regulating NF-κB-driven inflammatory responsiveness. Interestingly, this function is distinct from Integrator’s broader transcriptional role in RNAPII pause release since the endonuclease catalytic activity does not play a role in transcriptional responsiveness to TNF-α. This is demonstrated by performing a rescue experiment allowing the replacement of endogenous INTS11 with wild type or the catalytic dead (E203Q) which fully restored the TNF-α responsiveness (Fig. 1M, Fig. S1Q and R). Additionally, depletion of the non-catalytic subunit INTS6 likewise inhibited the TNF-α response (Fig. 1N, Fig. S1S) without destabilizing core catalytic subunits. Taken together, our results point to a critical role for Integrator in mediating the RELA transcriptional responsiveness following TNF-α or LPS stimulation in human and mouse cells, respectively.

### Integrator directs NF-κB association with enhancers at chromatin

We next used super-resolution nanoscopy (stochastic optical reconstruction microscopy [STORM]) to visualize the occupancy of RELA on chromatin fiber using antibodies against the core histone protein H3 at single cell level, with a resolution of ∼20 nm (Fig. 2A). While RELA is shown to associate with chromatin fiber (H3 colocalization) following TNF-α stimulation (Fig. 2A, depicted in Coloc-Tesseler analyses of colocalized image by red color), depletion of INTS11 specifically abrogates its interaction with chromatin without effecting its nuclear localization (Fig. 2A and B).

**Figure 2.**
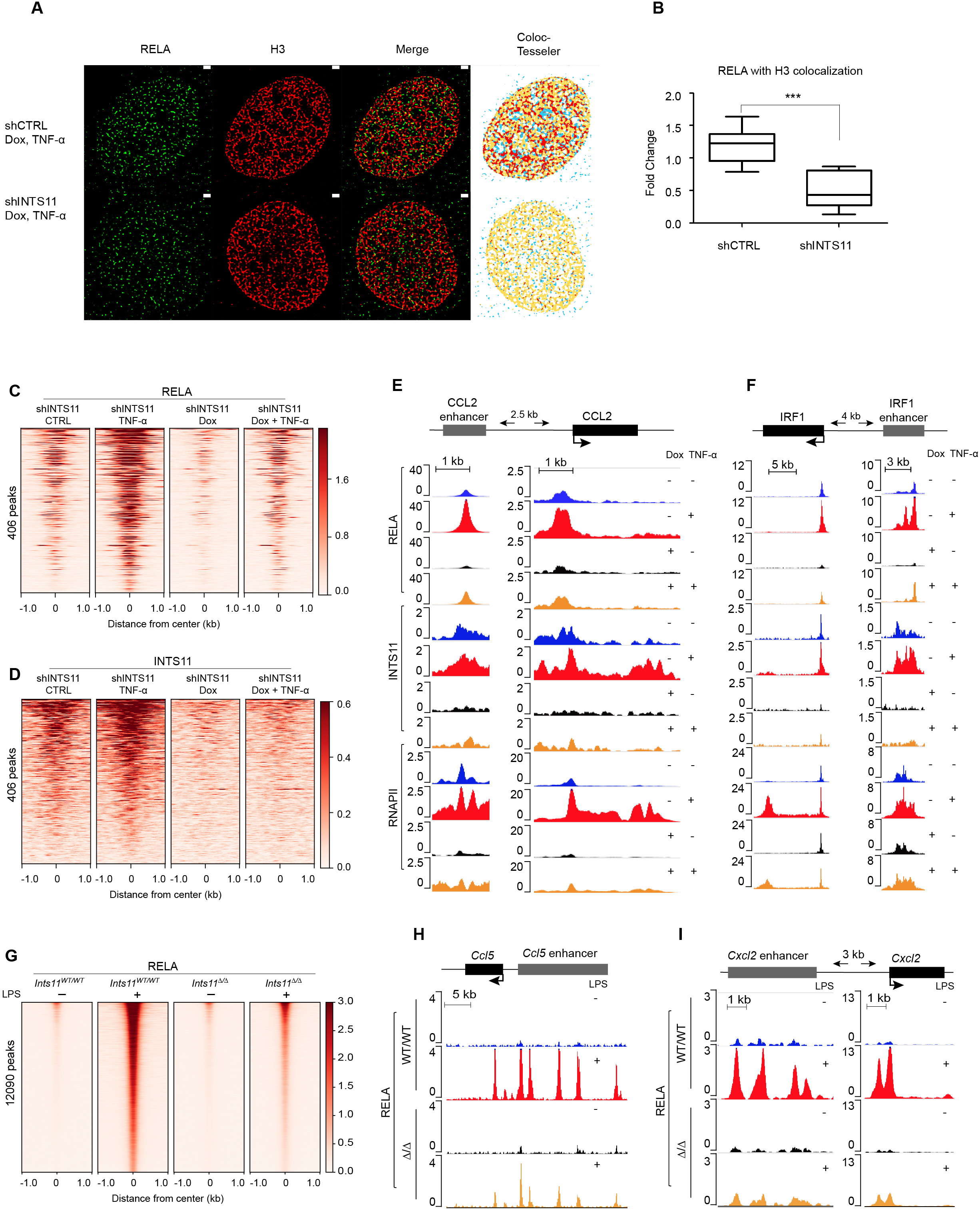
Integrator facilitates NF-κB association with enhancers on chromatin. **(A)** STORM imaging of RELA and histone H3 in Dox-treated shCTRL and shINTS11 HeLa cells following TNF-α stimulation. RELA and H3 signals, merged images, and spatial colocalization determined using Coloc-Tesseler are shown. **(B)** Quantification of RELA-H3 colocalization in Dox-treated, TNF-α–stimulated shCTRL and shINTS11 cells. ***P < 0.001 (n=10). **(C and D)** Heat maps showing RELA **(C)** and INTS11 **(D)** ChIP-seq occupancy centered on 406 RELA peaks at enhancers associated with 201 TNF-α–responsive genes in HeLa cells. Signals are shown in shINTS11 cells under control, TNF-α–stimulated, Dox-treated, and Dox plus TNF-α–treated conditions. **(E and F)** Representative ChIP-seq genome-browser tracks showing RELA, INTS11, and RNAPII occupancy at the *CCL2* gene and associated enhancer **(E)** and the *IRF1* gene and associated enhancer **(F)** in shINTS11 HeLa cells under the indicated Dox and TNF-α treatment conditions. Sequencing tracks were visualized in BigWig format using the WashU Epigenome Browser and aligned to the hg19 human genome assembly. **(G)** Heat maps showing RELA ChIP-seq occupancy centered on 12,090 RELA peaks in mouse bone marrow-derived macrophages (mBMDMs) from *Ints11^WT/WT^* and *Ints11*^Δ/Δ^ mice before and after LPS stimulation. **(H and I)** Representative RELA ChIP-seq genome-browser tracks at the *Ccl5* gene and associated enhancer **(H)** and the *Cxcl2* gene and associated enhancer **(I)** in mBMDMs from *Ints11^WT/WT^* and *Ints11*^Δ/Δ^ mice before and after LPS stimulation. Sequencing tracks were visualized in BigWig format using the WashU Epigenome Browser and aligned to the mm10 mouse genome assembly.

**Figure S2.**
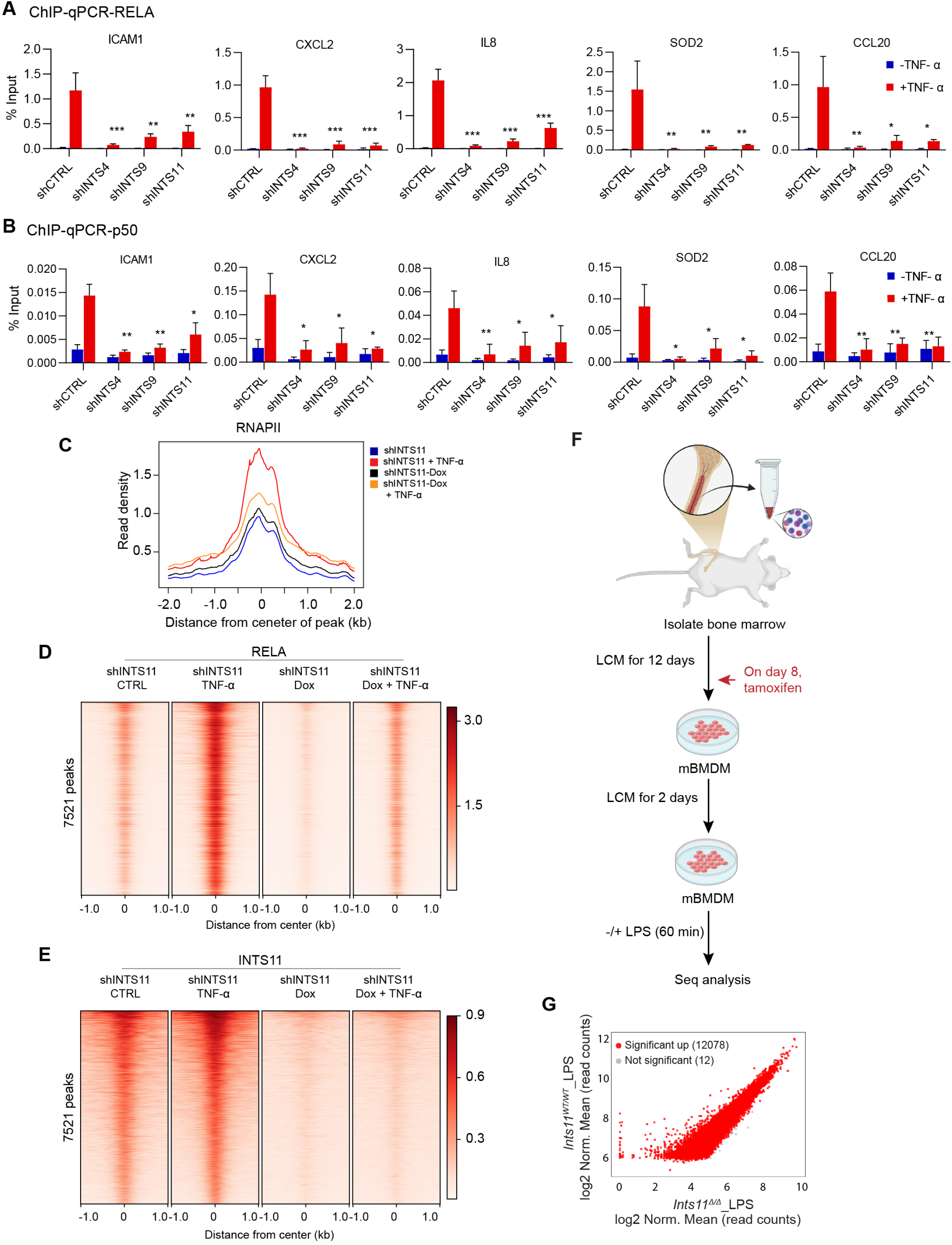
Integrator promotes NF-κB association with enhancers on chromatin. **(A and B)** ChIP-qPCR analysis of RELA **(A)** and p50 **(B)** occupancy at the TNF-α–responsive *ICAM1*, *CXCL2*, *IL8*, *SOD2*, and *CCL20* loci in HeLa shCTRL, shINTS4, shINTS9, and shINTS11 cells before and after TNF-α stimulation. ChIP enrichment is expressed as percentage of input. Error bars represent SD from at least three independent biological experiments (*n* ≥ 3). Statistical significance was determined using an unpaired two-tailed Student’s *t* test. *P < 0.05; **P < 0.01; ***P < 0.001. **(C)** Metaplots profiles of RNAPII ChIP-seq occupancy centered on RELA peaks at enhancers associated with 201 TNF-α–responsive genes in shINTS11 HeLa cells under the indicated Dox and TNF-α treatment conditions. **(D and E)** Heat maps showing genome-wide RELA **(D)** and INTS11 **(E)** ChIP-seq occupancy centered on 7,521 RELA peaks associated with enhancers in shINTS11 HeLa cells under the indicated Dox and TNF-α treatment conditions. **(F)** Experimental scheme for generation and analysis of mouse bone marrow-derived macrophages (mBMDMs). Bone marrow cells were cultured in L929-conditioned medium (LCM), with tamoxifen treatment initiated on day 8 as indicated. Following differentiation, mBMDMs were cultured further in LCM and subsequently left unstimulated or stimulated with LPS for 60 min before sequencing analysis. **(G)** Scatter plot comparing RELA ChIP-seq occupancy at RELA-associated chromatin regions in LPS-stimulated mBMDMs from *Ints11^WT/WT^* and *Ints11*^Δ/Δ^ mice. Axes indicate log2-normalized mean RELA ChIP-seq read counts. Differentially occupied regions are indicated in red (12,078), and regions without a significant difference are shown in gray (12).

**Figure S3.**
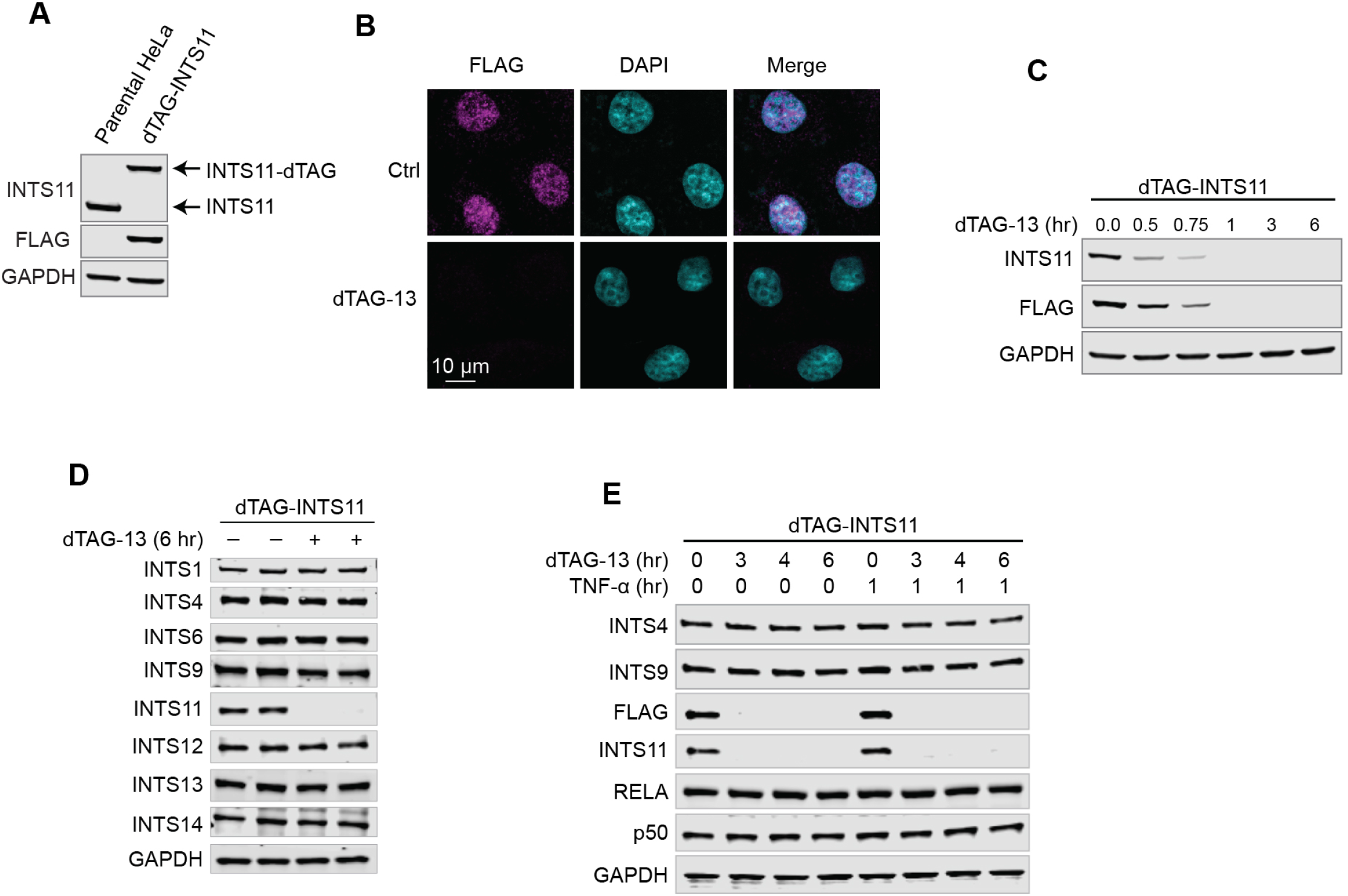
Characterization of the dTAG-INTS11 acute-degradation system. **(A)** Immunoblot analysis of parental HeLa and dTAG-INTS11 cells showing endogenous INTS11 and the higher-molecular-weight dTAG-tagged INTS11 protein. FLAG confirms expression of the tagged INTS11 protein, and GAPDH served as a loading control. **(B)** Representative immunofluorescence images showing FLAG-tagged dTAG-INTS11 in control and dTAG-13–treated cells. DAPI marks nuclei. Scale bar, 10 μm. **(C)** Time-course analysis of dTAG-INTS11 degradation following addition of dTAG-13 for 0, 0.5, 0.75, 1, 3, or 6 h. INTS11 and FLAG were detected by immunoblotting, with GAPDH as a loading control. **(D)** Immunoblot analysis of Integrator subunits following acute INTS11 degradation. INTS1, INTS4, INTS6, INTS9, INTS11, INTS12, INTS13, and INTS14 were examined in dTAG-INTS11 cells treated with or without dTAG-13 for 6 h. GAPDH served as a loading control. **(E)** Immunoblot analysis of INTS4, INTS9, FLAG-tagged INTS11, endogenous INTS11, RELA, and p50 during a dTAG-13 time course in the absence or presence of TNF-α stimulation. Cells were exposed to dTAG-13 for 0, 3, 4, or 6 h and were either left unstimulated or stimulated with TNF-α for 1 h before collection. GAPDH served as a loading control.

To assess changes in NF-κB occupancy at enhancers of inflammatory genes, we performed chromatin immunoprecipitation followed by sequencing (ChIP-seq) using RELA, INTS11 and RNAPII antibodies prior to and following stimulation of HeLa cells with TNF-α. It is important to note that NF-κB is known to be consecutively active in many cancers including cervical cancer(*26, 27*). We initially assessed RELA occupancy at enhancers within a 20 kilobase window in the proximity of the transcription start sites of TNF-α responsive genes (406 RELA peaks) (Fig. 2C-F, Table 3). Consistent with the diminished transcription of inflammatory genes (Fig. 1) and observed loss of RELA association with chromatin using STORM analyses, INTS11 depletion leads to a marked decrease in RELA association at enhancers of TNF-α responsive genes (Fig. 2C-F). We validated these results using chromatin immunoprecipitation followed by real-time PCR demonstrating decreased association of RELA and its partner p50 following depletion of INTS11, INTS4 and INTS9 at enhancers of five TNF-α -induced genes in HeLa cells (Fig. S2A and B). Additionally, consistent with the loss of transcriptional responsiveness, INTS11 depletion leads to a decreased RNAPII recruitment at regulatory sites (Fig. 2E and F, and Fig. S2C). Indeed, genome-wide analyses of RELA sites that are co-occupied by INTS11 (7521 peaks) revealed loss of NF-κB binding upon depletion of INTS11 (Fig. S2D and E, Table 3). To extend these findings, we assess RELA association with chromatin in bone-marrow derived macrophages following stimulation with LPS in wild type mice and mice with deletion of *Ints11* (Fig. S2F). We find that deletion of INTS11 result in a highly significant loss of RELA association with chromatin genome-wide following LPS induction (12,078 sites loss RELA association significantly, Fig. 2G-I, Fig. S2G, Table 4). Taken together, these results highlight a critical role for Integrator in promoting NF-κB binding to chromatin at inflammatory genes in human and mouse cells following an inflammatory stimulus.

### Acute depletion of INTS11 leads to loss of NF-κB association with enhancers

We developed a degron system for INTS11 in HeLa cells using dTAG-INTS11 knock-in which displaying a similar localization as endogenous INTS11 (Fig. 3A, Fig. S3A-B). Using this system INTS11 could be depleted within 1 hour of dTAG-13 addition (Fig. S3C). Depletion of dTAG-INTS11 does not affect the levels of other Integrator subunits or components of NF-κB within the 6-hour time point (Fig. S3D and E). Importantly, a 6-hour depletion of dTAG-INTS11 does not affect the TNF-α-induced translocation of RELA into the nucleus or the degradation of IkBα, confirming that acute depletion of INTS11 does not disrupt the cytoplasmic to nuclear translocation of NF-κB (Fig. 3B-F). Finally, single molecule imaging confirms the colocalization of d-TAG-INTS11 and RELA using super-resolution microscopy (Fig. 3G).

**Figure 3.**
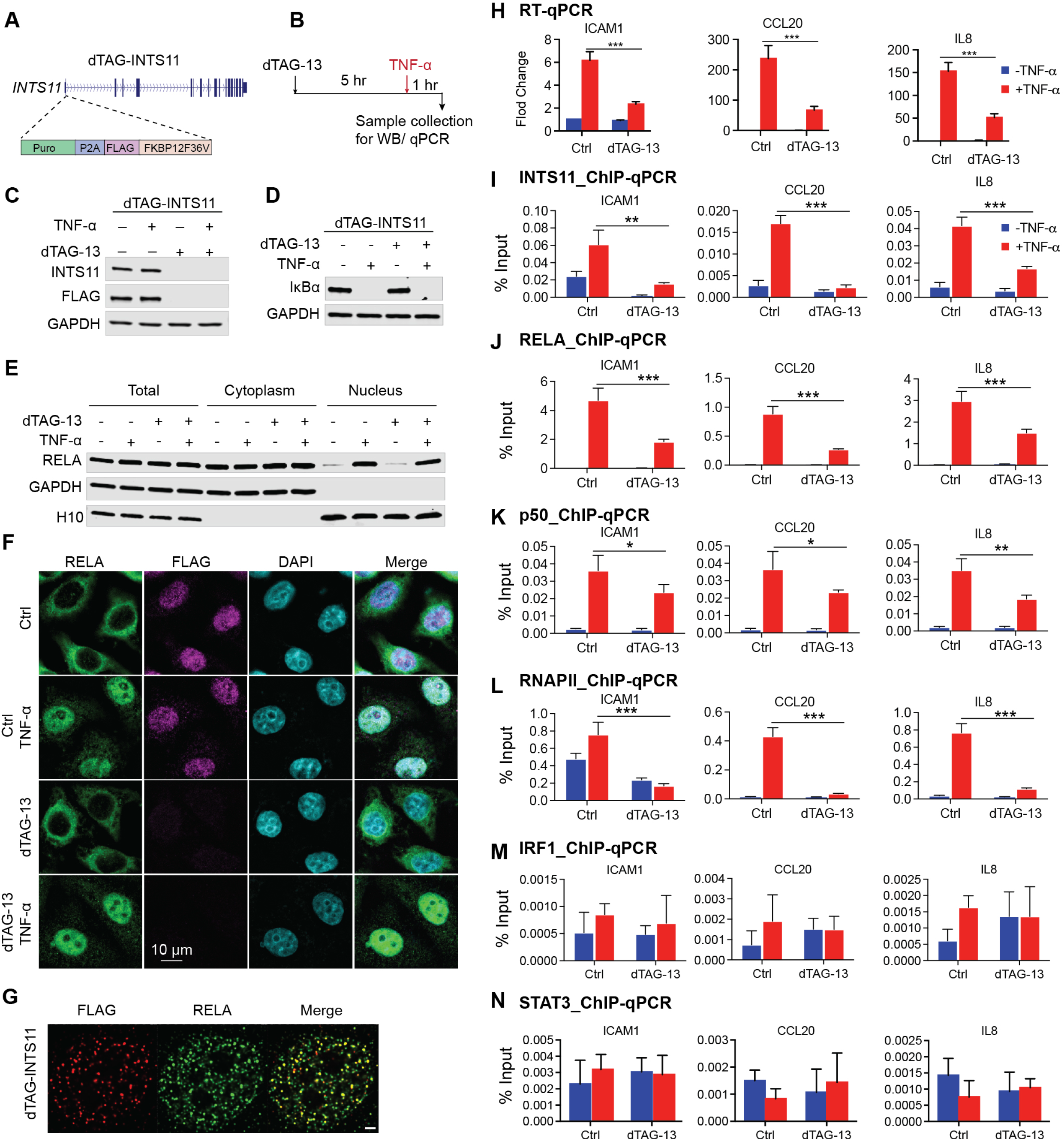
Acute depletion of INTS11 impairs NF-κB recruitment and transcriptional activation at inflammatory genes. **(A)** Schematic of the endogenous dTAG-INTS11 allele generated in HeLa cells. The C terminus of INTS11 was fused to a FLAG-tagged FKBP12^F36V^ degron cassette. **(B)** Experimental scheme for acute INTS11 degradation and TNF-α stimulation. dTAG-INTS11 cells were treated with dTAG-13 for 5 h, followed by TNF-α stimulation for 1 h, for a total of 6 h of dTAG-13 treatment before collection for immunoblotting or qPCR analysis. **(C)** Immunoblot analysis of INTS11 degradation in dTAG-INTS11 cells treated with dTAG-13 and/or TNF-α as indicated. INTS11 and FLAG were detected, and GAPDH served as a loading control. **(D)** Immunoblot analysis of IκBα in dTAG-INTS11 cells treated with dTAG-13 and/or TNF-α. GAPDH served as a loading control. **(E)** Subcellular fractionation and immunoblot analysis of RELA in dTAG-INTS11 cells treated with dTAG-13 and/or TNF-α. RELA abundance is shown in total, cytoplasmic, and nuclear fractions. GAPDH and histone H1.0 served as cytoplasmic and nuclear fraction markers, respectively. **(F)** Representative immunofluorescence images showing RELA, FLAG-tagged dTAG-INTS11, and DAPI in control and dTAG-13–treated cells before and after TNF-α stimulation. Scale bar, 10 μm. **(G)** Representative STORM images showing FLAG-tagged dTAG-INTS11 and RELA and their merged localization following TNF-α stimulation. **(H)** RT-qPCR analysis of the TNF-α–responsive genes *ICAM1*, *CCL20*, and *IL8* in control and dTAG-13–treated cells before and after TNF-α stimulation. **(I to N)** ChIP-qPCR analysis of factor occupancy at the *ICAM1*, *CCL20*, and *IL8* loci in control and dTAG-13–treated cells before and after TNF-α stimulation. Occupancy of INTS11 **(I)**, RELA **(J)**, p50 **(K)**, RNAPII **(L)**, IRF1 **(M)**, and STAT3 **(N)** was measured and expressed as percentage of input. For **(H to N)**, error bars represent SD from at least three independent biological experiments (*n* ≥ 3). Statistical significance was determined using an unpaired two-tailed Student’s *t* test for the indicated comparisons. *P* < 0.05; **P < 0.01; ***P < 0.001.

We next assessed whether acute depletion of INST11 impacts NF-κB (RELA and p50) as well as RNAPII recruitment to a set of key TNF-α responsive genes using ChIP-qPCR and RT-qPCR (Fig. 3H-L). Importantly, ICAM1, CCL20, IL8 display detectable levels of INTS11 occupancy prior to TNF-α stimulation in the absence of any significant NF-κB residence, however, 1 hour following TNF-α induction, there is a robust recruitment of INTS11 and NF-κB to all three genes (Fig. 3I-K). Notably, acute depletion of INTS11 (6 hours), significantly diminish NF-κB recruitment concomitant with the loss of RNAPII deployment and induction of transcription (Fig. 3H-L). The INTS11-mediated recruitment of NF-κB at these promoters is specific since we don’t observe a significant change in occupancy of other transcription factors occupying these promoters such as IRF1 or STAT3 (Fig. 3M and N). These results reveal the crucial role for INST11 in recruitment of NF-κB to inflammatory genes and pinpoints a direct role for Integrator in induction of inflammatory transcriptional response.

### Integrator controls NF-κB residence times using single molecule tracking

To gain insight into the mechanism by which INTS11regulates RELA association with chromatin, we performed single-molecule tracking (SMT) experiments using a Halo-tag RELA following acute depletion of INTS11 (Fig. 4A-D). The three populations of molecules detected by SMT (Fig. 4E and F, unbound, fast bound and slow bound fractions) has been interpreted as the transcription factors diffusing in the nucleus (unbound fraction), binding to non-specific sites in chromatin (fast component), or to specific response elements (slow component)(*28*). Interestingly, while acute depletion of INTS11 does not significantly change the percentage of each fraction, we observe a profound decrease in the residence time for both the slow and fast fractions (Fig. 4E and F), reflective of INTS11 requirement for RELA association with its cognate sites *in vivo*. Indeed, the slow fraction representing the specific binding events for RELA to its cognate sites decreased from about 4 seconds to 1.4 seconds, highlighting the importance of INTS11 for specific association of NF-κB to chromatin (Fig. 4G). Importantly, while we observe a similar trend after depletion of INTS11 using shRNAs against INTS11, replacing wild type INTS11 with its catalytic mutant E203Q did not affect RELA binding residence times (Fig. 4H and I-M), reinforcing the contention that the catalytic activity is not critical for the residence time.

**Figure 4.**
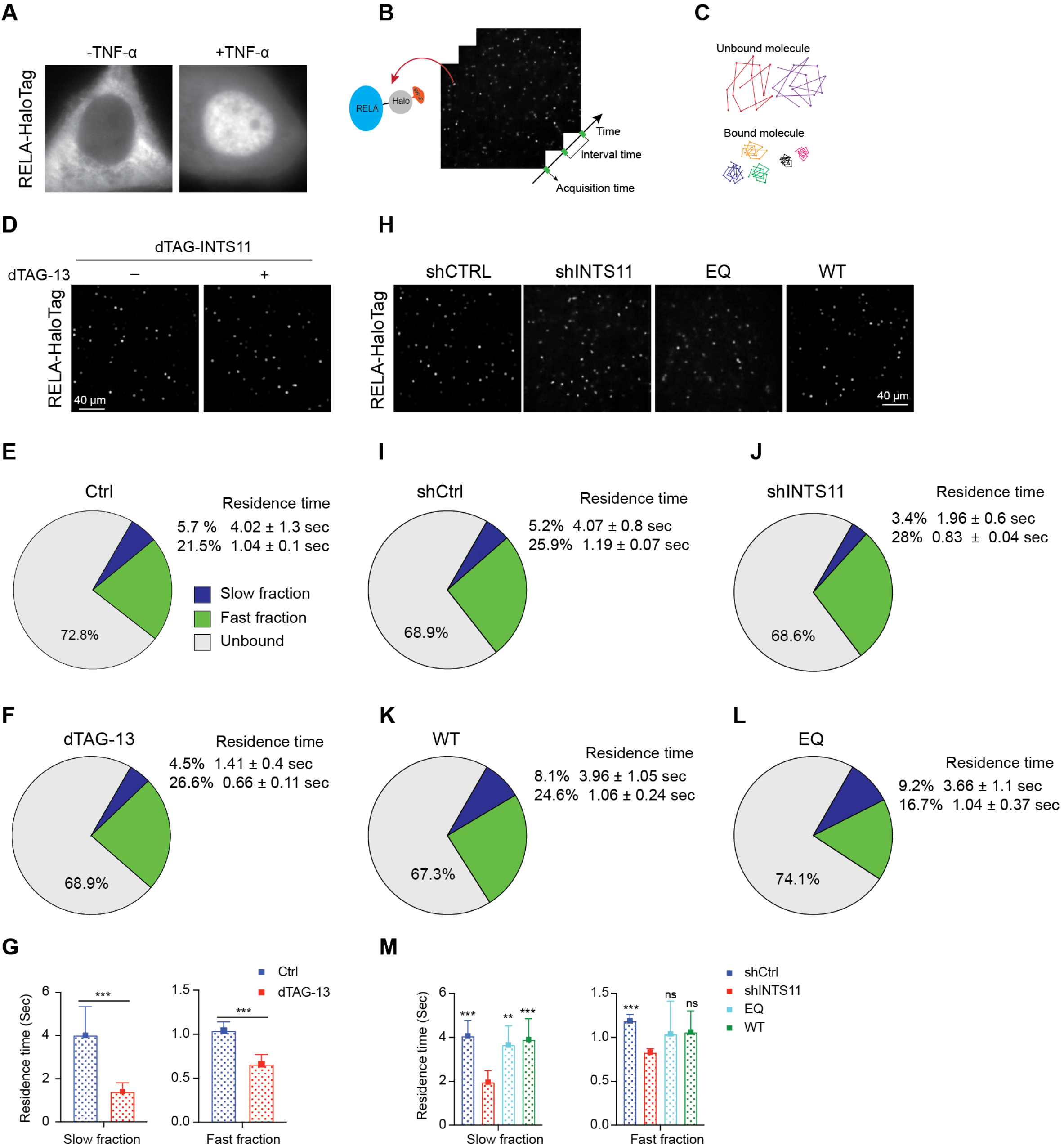
Integrator controls NF-κB chromatin residence time. **(A)** Fluorescence images showing RELA-HaloTag localization before and after TNF-α stimulation. **(B)** Schematic of the single-molecule tracking (SMT) approach used to measure RELA-HaloTag dynamics in living cells. RELA-HaloTag was fluorescently labeled and individual molecules were imaged over time at defined acquisition intervals. **(C)** Representative trajectories illustrating the classification of RELA molecules as bound or unbound. Bound molecules exhibit spatially confined trajectories, whereas unbound molecules display greater diffusive movement. **(D)** Representative TIRF-based single-molecule images of RELA-HaloTag in dTAG-INTS11 cells treated with or without dTAG-13. Scale bar, 40 μm. **(E and F)** Fractions of unbound, slow-fraction, and fast-fraction RELA-HaloTag molecules in control **(E)** and dTAG-13–treated **(F)** cells. The corresponding residence times of the slow- and fast-fraction fractions are indicated. **(G)** Quantification of residence times for the slow- and fast-fraction RELA-HaloTag molecules in control and dTAG-13–treated cells. Data are presented as mean ± SD. Statistical significance was determined using Student’s *t* test for the indicated comparisons. ***P < 0.001 (n≥10). **(H)** Representative TIRF-based single-molecule images of RELA-HaloTag in shCTRL and shINTS11 cells and in INTS11-depleted cells re-expressing E203Q-INTS11 or WT-INTS11 under the indicated treatment conditions. Scale bar, 40 μm. **(I to L)** Fractions of unbound, slow-fraction, and fast-fraction RELA-HaloTag molecules in shCTRL **(I)**, shINTS11 **(J)**, WT-INTS11–reconstituted **(K)**, and E203Q-INTS11–reconstituted **(L)** cells. Corresponding residence times of the slow- and fast-fractions are indicated. **(M)** Quantification of residence times for the slow- and fast-fraction RELA-HaloTag molecules in shCTRL and shINTS11 cells and in INTS11-depleted cells re-expressing E203Q-INTS11 or WT-INTS11. Data represent mean ± SD. Statistical differences were evaluated by one-way ANOVA followed by Dunnett’s multiple comparisons test against the shINTS11 condition (**p < 0.01, ***p < 0.001). (n≥10).

**Figure S4.**
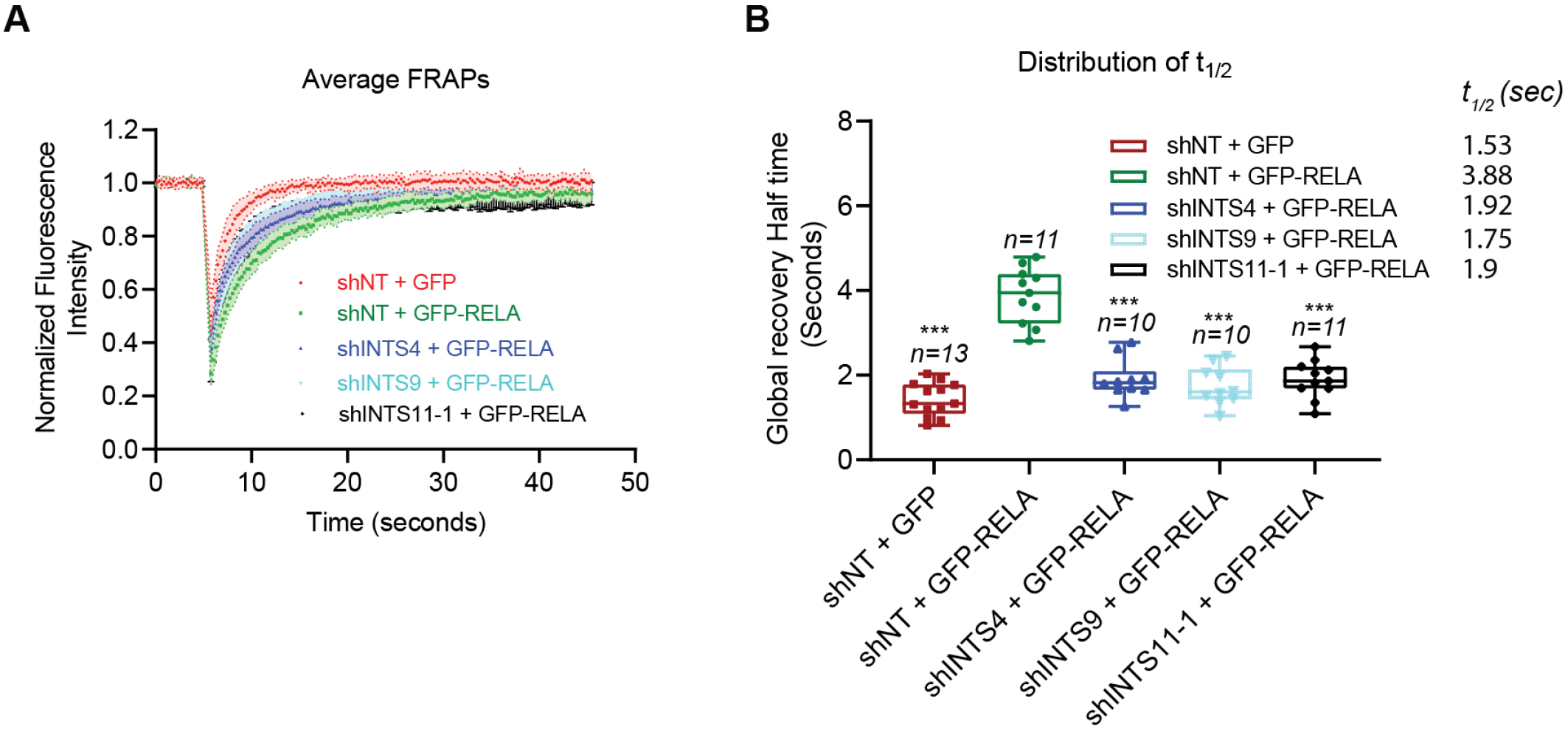
Integrator regulates RELA nuclear mobility. **(A)** Average fluorescence recovery after photobleaching (FRAP) curves for GFP and GFP-RELA in control cells and for GFP-RELA following depletion of INTS4, INTS9, or INTS11. Fluorescence intensity was normalized to the prebleach signal. **(B)** Distribution of global fluorescence recovery half-times (*t*_1/2_ calculated from the FRAP experiments in **(A)**. Mean *t*_1/2_ values were 1.53 s for shNT + GFP, 3.88 s for shNT + GFP-RELA, 1.92 s for shINTS4 + GFP-RELA, 1.75 s for shINTS9 + GFP-RELA, and 1.90 s for shINTS11 + GFP-RELA. Sample sizes are indicated in the plot. ***P < 0.001 for the indicated comparisons.

To independently assess RELA dynamics, we next used fluorescence recovery after photobleaching (FRAP) to further assess the nuclear mobility of RELA and its dependence on Integrator following induction with TNF-α (Fig. S4A and B). While RELA displayed a global recovery half-time of approximately 4 seconds following induction with TNF-α, depletion of INTS4, INTS9 or INTS11 led to a decrease of nuclear mobility to below a half-time of 2 seconds (Fig. S4A and B). These results confirm that the core Integrator catalytic module is critical for RELA mobility through the chromatin environment, manifested as a shorter half-life in the absence of Integrator and indicative of decreased chromatin association *in vivo*. Collectively, Integrator acts as a direct molecular anchor, stabilizing RELA’s binding affinity and increasing its chromatin residence time, a function that is independent of its catalytic activity. This mechanism licenses the stable NF-κB association necessary for transcriptional output *in vivo*, mirroring the results obtained using chromatin immunoprecipitation.

### NF-κB associates with Integrator on chromatin

To determine whether Integrator interacts with NF-κB, we performed an immunoprecipitation using INTS11 antibodies following TNF-α stimulation (Fig. 5A). We find that Integrator interaction with NF-κB is DNA dependent since treatment of immunoprecipitated eluate with DNase I substantially diminished their association (Fig. 5B). We next assessed the proximity of Integrator and NF-κB binding sites at enhancers on chromatin by reChIP or ChIP-on-chip experiments where INTS11 ChIP was used to perform a RELA ChIP-seq. As Figures 5C-E demonstrates INTS11 binding sites at enhancers are highly enriched with RELA occupancy. The proximity of INTS11 and RELA binding on chromatin is seen genome-wide at 7521 co-occupied sites (Fig. S5A). Finally, replacement of wild type INTS11 with a catalytically dead mutant does not affect TNF-α−dependent recruitment of RELA highlighting a catalytic-independent function of Integrator in mediating NF-κB association with chromatin (Fig. 5F-H and Fig. S5C-D). We find less than 10% of RELA binding is significantly affected by replacement of INTS11 with a catalytically dead mutant (Fig. S5F). These results support the contention that Integrator and NF-κB localize to the same sites on chromatin and the two protein complexes are in close proximity on enhancers of inflammatory genes.

**Figure 5.**
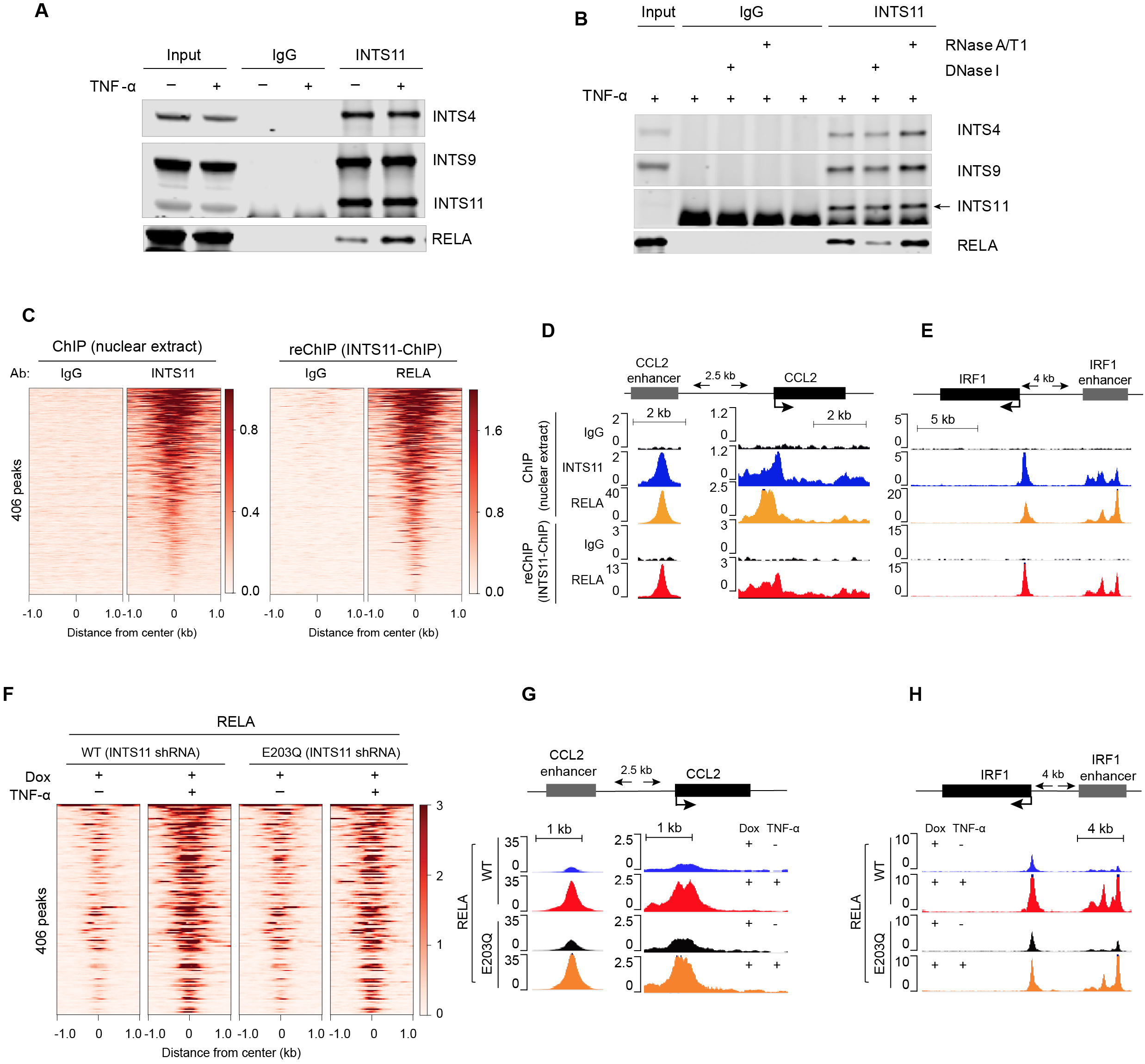
NF-κB associates with Integrator on chromatin. **(A)** Co-immunoprecipitation analysis of endogenous INTS11 in HeLa cells before and after TNF-α stimulation. INTS11 was immunoprecipitated, and associated INTS4, INTS9, INTS11, and RELA were detected by immunoblotting. **(B)** Co-immunoprecipitation analysis of the INTS11–RELA association following nuclease treatment TNF-α-stimulated HeLa cell extracts. HeLa cell extracts were immunoprecipitated with INTS11 or control IgG antibodies and treated with DNase I or RNase A/T1, as indicated. INTS4, INTS9, INTS11, and RELA were detected by immunoblotting. **(C)** Heat maps showing INTS11 ChIP-seq and sequential INTS11–RELA reChIP signals centered on 406 RELA peaks at enhancers associated with 201 TNF-α–responsive genes in HeLa cells. For reChIP, chromatin initially immunoprecipitated with INTS11 antibody was subjected to a second immunoprecipitation with RELA or control IgG antibody. **(D and E)** Representative genome-browser tracks showing INTS11 and RELA ChIP-seq occupancy and sequential INTS11–RELA reChIP signal at the *CCL2* gene and proximal enhancer **(D)** and the *IRF1* gene and proximal enhancer **(E)** following TNF-α stimulation in HeLa cells. **(F)** Heat maps showing RELA ChIP-seq occupancy centered on the 406 RELA peaks at enhancers associated with 201 TNF-α–responsive genes in INTS11-depleted cells re-expressing WT-INTS11 or the E203Q-INTS11 catalytic mutant, before and after TNF-α stimulation. **(G and H)** Representative genome-browser tracks showing RELA ChIP-seq occupancy at the *CCL2* gene and proximal enhancer **(G)** and the *IRF1* gene and proximal enhancer **(H)** in INTS11-depleted HeLa cells re-expressing WT-INTS11 or E203Q-INTS11, before and after TNF-α stimulation.

**Figure S5.**
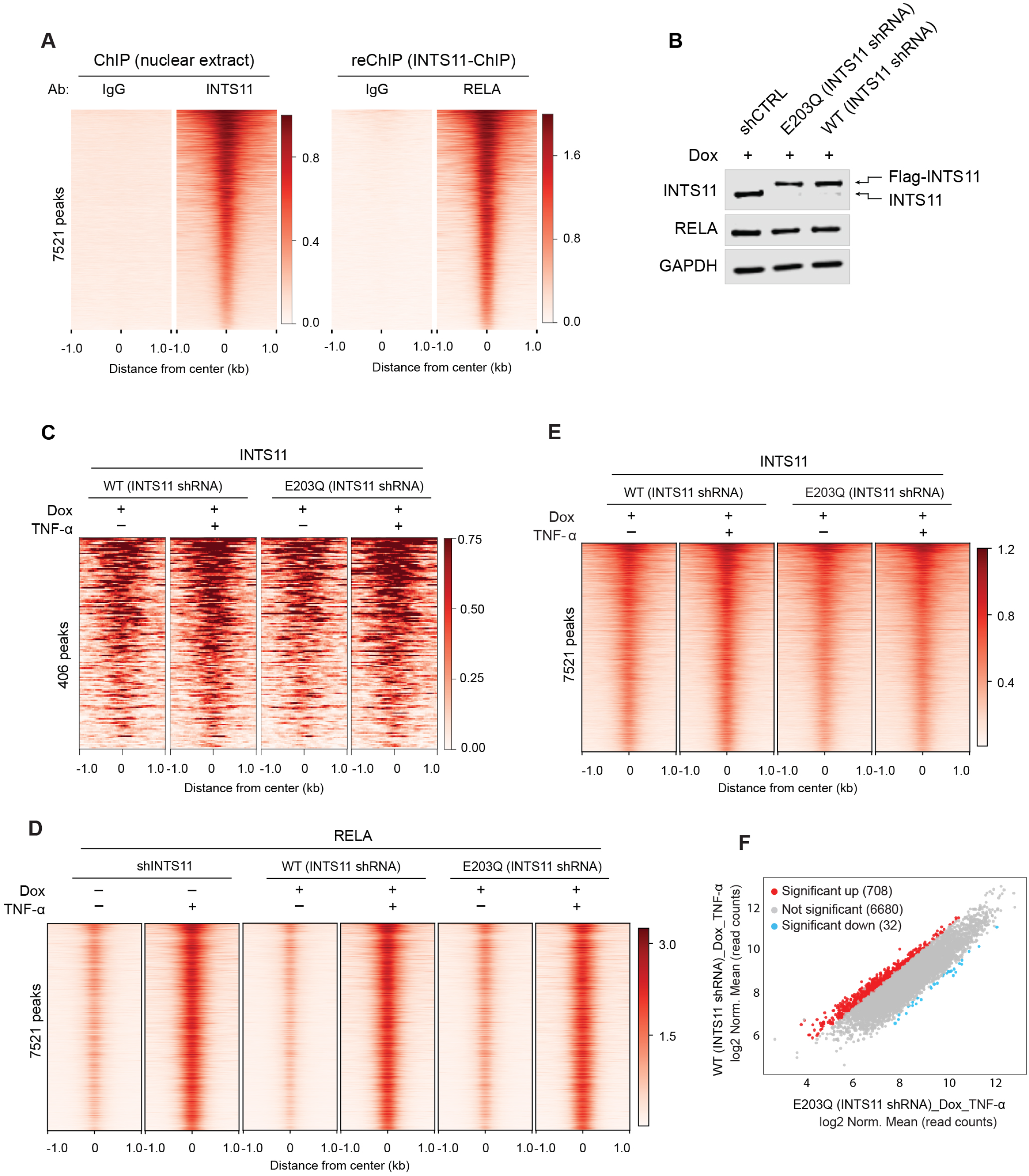
Genome-wide association of NF-κB and Integrator on chromatin. **(A)** Heat maps showing genome-wide INTS11 ChIP-seq and sequential INTS11–RELA reChIP signals centered on 7,521 RELA peaks associated with enhancers. For reChIP, chromatin initially immunoprecipitated with INTS11 antibody was subjected to a second immunoprecipitation with RELA or control IgG antibody. **(B)** Immunoblot analysis of INTS11 shRNA cells reconstituted with WT-INTS11 or E203Q-INTS11. FLAG-tagged reconstituted INTS11 and endogenous INTS11 are indicated; RELA levels are also shown. GAPDH served as a loading control. **(C)** Heat maps showing INTS11 ChIP-seq occupancy centered on the 406 RELA peaks at enhancers associated with 201 TNF-α–responsive genes **in** INTS11 shRNA cells reconstituted with WT-INTS11 or E203Q-INTS11, before and after TNF-α stimulation. **(D)** Heat maps showing genome-wide RELA ChIP-seq occupancy centered on 7,521 RELA peaks associated with enhancers in INTS11 shRNA cells reconstituted with WT-INTS11 or E203Q-INTS11, before and after TNF-α stimulation. **(E)** Heat maps showing genome-wide INTS11 ChIP-seq occupancy centered on the same 7,521 RELA peaks associated with enhancers **in** INTS11 shRNA cells reconstituted with WT-INTS11 or E203Q-INTS11, before and after TNF-α stimulation. **(F)** Scatter plot comparing RELA ChIP-seq occupancy in TNF-α–stimulated INTS11 shRNA HeLa cells reconstituted with WT-INTS11 or E203Q-INTS11.

### Whole body deletion of *Ints11* in adult mice affects the hematopoietic system

We developed an inducible whole body knockout mouse model for *Ints11* to gain insight into its biological function by crossing *Ints11^Flox/Flox^*with *Rosa26-Cre-ERT2^+^* mice. To access the consequences of whole body INTS11 deletion, young adult mice (aged 2-4 months) with correct genotypes were injected intraperitoneally with tamoxifen (TAM) at a dosage of 100 mg/kg once per day for five consecutive days to delete *Ints11* and survival was monitored. The survival curve shows that the INTS11-WT mice (*Rosa26-Cre-ERT2^+^*) didn’t display a significant effect from TAM injection, whereas the INTS11-KO mice (*Ints11^flox/flox^; Rosa26-Cre-ERT2^+^*) exhibited significantly reduced body weight during the TAM treatment leading to their death within 2 weeks of TAM treatment with a median survival of 8.5 days (Fig. 6A-B). Consistent with deletion of *Ints11*, tissues collected for RT-qPCR analyses 7 days post-TAM injection demonstrated significantly lower mRNA levels in the bone marrow, spleen, lung, liver, and kidney of INTS11-KO mice compared to INTS11-WT mice (Fig. 6C). These results indicate that the loss of INTS11 in adult mice leads to a marked decrease in animals fitness leading to the loss of survival in less than two weeks.

**Figure 6.**
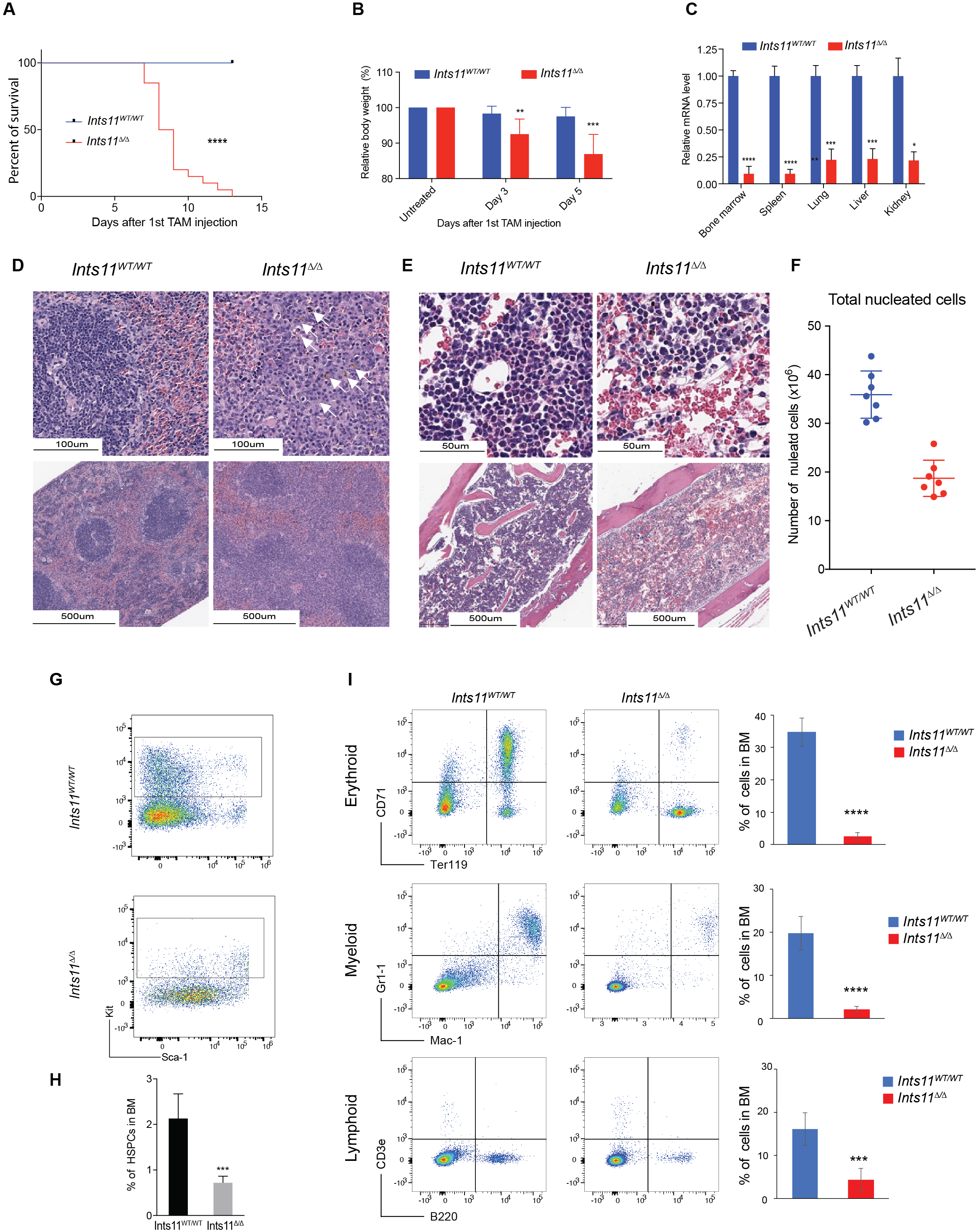
Whole-body deletion of *Ints11* compromises survival and hematopoiesis in adult mice. **(A)** Kaplan-Meier survival analysis of *Ints11^WT/WT^* and *Ints11*^Δ/Δ^ mice following tamoxifen (TAM) administration. Mice 2 to 4 months of age received TAM (100 mg/kg) once daily for five consecutive days and were monitored for survival. *n* = 20 mice per genotype. Statistical significance was determined using the log-rank (Mantel-Cox) test; ****P < 0.0001. **(B)** Relative body weight of *Ints11^WT/WT^* and *Ints11*^Δ/Δ^ mice before TAM treatment and at days 3 and 5 after the first TAM injection. Values are expressed relative to pretreatment body weight. **(C)** Relative *Ints11* mRNA levels in bone marrow, spleen, lung, liver, and kidney collected 7 days after TAM administration from *Ints11^WT/WT^* and *Ints11*^Δ/Δ^ mice. *Ints11* transcript abundance was measured by RT-qPCR and expressed relative to the corresponding *Ints11^WT/WT^* control. Data represent three biological replicates per group. Statistical significance was determined using Student’s *t* test. **(D)** Representative hematoxylin and eosin (H&E)-stained spleen sections from *Ints11^WT/WT^* and *Ints11*^Δ/Δ^ mice collected 7 days after TAM administration. Upper and lower panels show higher- and lower-magnification views, respectively. White arrowheads indicate increased pigment accumulation in the *Ints11*^Δ*/*Δ^ spleen. Scale bars, 100 μm (top) and 500 μm (bottom). **(E)** Representative H&E-stained bone marrow sections from *Ints11^WT/WT^* and *Ints11*^Δ/Δ^ mice collected 7 days after TAM administration. Upper and lower panels show higher- and lower-magnification views, respectively. Scale bars, 50 μm (top) and 500 μm (bottom). **(F)** Number of total viable nucleated cells in bone marrow. **(G)** Representative flow-cytometry plots showing the c-Kit^+^ hematopoietic stem/progenitor cell (HSPC) population when gated on Lin− live bone marrow cells from *Ints11^WT/WT^* and *Ints11*^Δ/Δ^ mice. **(H)** Quantification of HSPCs (Lin^−^ c-Kit^+^ population) in bone marrow from *Ints11^WT/WT^* and *Ints11*^Δ/Δ^ mice, expressed as a percentage of analyzed bone marrow cells. **(I)** Representative flow-cytometry plots and corresponding quantification of erythroid, myeloid, and lymphoid populations in bone marrow from *Ints11^WT/WT^* and *Ints11*^Δ/Δ^ mice. Erythroid cells were identified as CD71+Ter119+, myeloid cells as Gr-1+Mac-1+, and B-lymphoid cells as CD3ε−B220+. Bar graphs show the percentage of each population among analyzed bone marrow cells. For **(B), (C), (F), (H), and (I)**, error bars represent SD. Statistical significance is indicated as *P < 0.05; **P < 0.01; ***P < 0.001; ****P < 0.0001.

**Figure S6.**
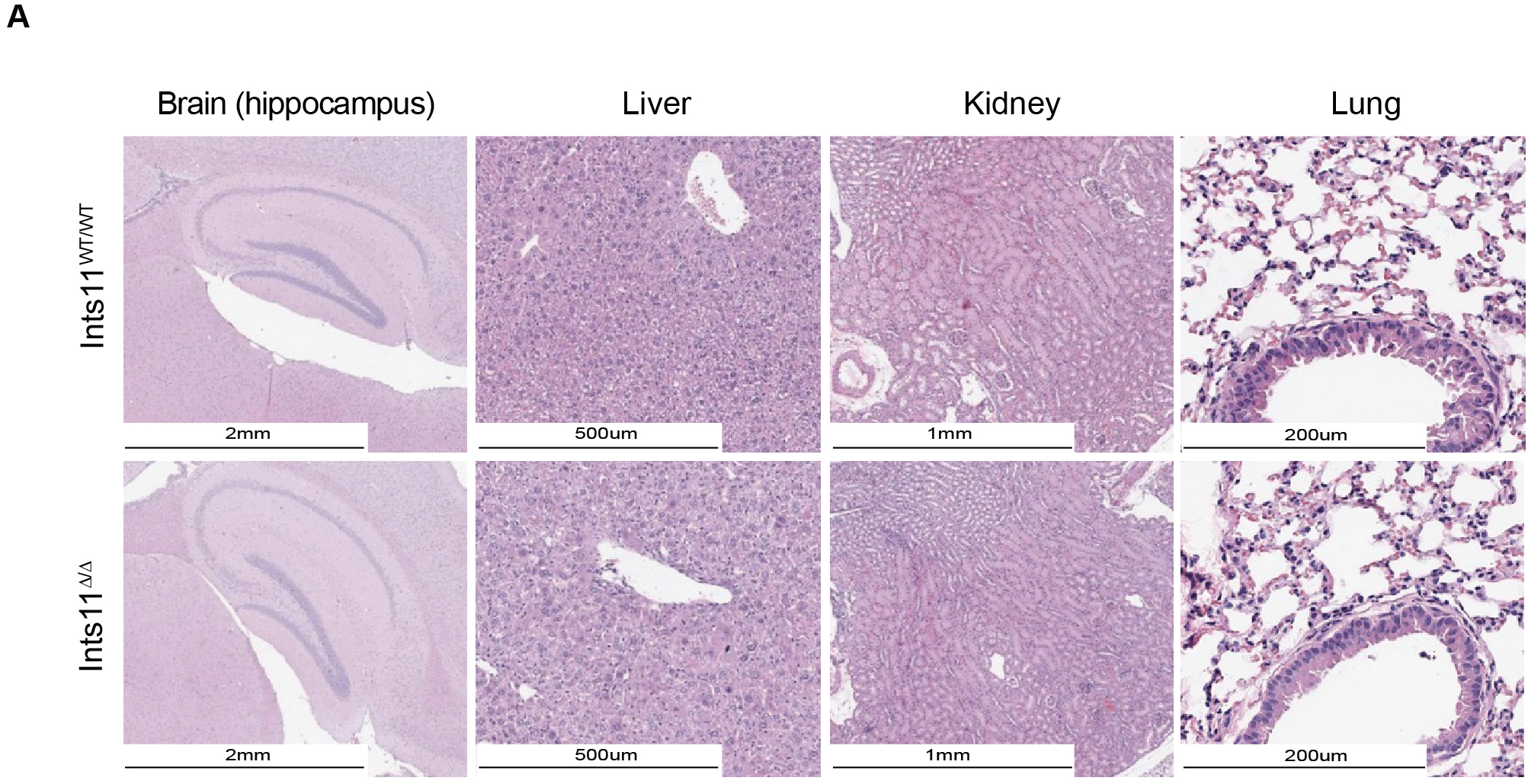
Histopathological evaluation of additional tissues following whole-body *Ints11* deletion. **(A)** Representative H&E-stained sections of brain (hippocampus), liver, kidney, and lung from *Ints11^WT/WT^* and *Ints11*^Δ/Δ^ mice collected 7 days after TAM administration. Corresponding tissues from the two genotypes are shown for comparison. Scale bars: brain (hippocampus), 2 mm; liver, 500 μm; kidney, 1 mm; and lung, 200 μm.

To gain insight into INTS11 loss of function in different tissues of adult mice, we evaluated the status of various organs, with a particular focus on the spleen, liver, brain, kidney, lung, and bone marrow, using hematoxylin and eosin (H&E) staining 7 days after tamoxifen injection. Comparative pathological evaluations between *Ints11*^WT/WT^ and *Ints11*^Δ/Δ^ groups were conducted to identify any morphological changes. We did not find any significant morphological changes in the brain, liver, kidney, or the lung between the two groups, as seen in the comparative histological images (Fig. S6A). In contrast, analyses of spleen displayed decreased megakaryocytes, disorganized white pulps and increased pigment infiltration (indicated by white arrows) in *Ints11*^Δ*/*Δ^ group (Fig. 6D). In addition, analyses of bone marrow (BM) revealed diminished cellularity and decreased megakaryocytes in *Ints11*^Δ*/*Δ^ group as compared to *Ints11^WT/WT^* (Fig. 6E). The count of total viable nucleated cells also supported this notion (Fig. 6F). Taken together, while the knockout of INTS11 in adult mice did not lead to a significant morphological change in most organs, alterations in spleen cellularity and bone marrow composition highlight its critical role in hematopoiesis and immune cell function.

To further dissect INTS11’s function in adult murine immune system, we sought to delineate the changes in cellular composition of bone marrow and spleen using flow cytometry. Interestingly, several recent studies have demonstrated that the steady-state hematopoiesis in normal adult mice is maintained by a large pool of self-renewing progenitor cells(*29*). Importantly, our flow cytometry analyses of bone marrow cells revealed a significant reduction in HSPC (Lin^−^ c-Kit^+^) representing the hematopoietic stem/progenitor cells population in INTS11-KO mice as compared to WT controls 7 days following TAM treatment (Fig. 6G and H). Additionally, the populations of erythroid, myeloid and lymphoid lineages in BM were significant reduced (Fig. 6I), reflecting a major bone marrow failure resulting from INTS11 deletion. Taken togther, these results reflects a profund role for INTS11 in early dvelopmental process of hematopoietic lineage.

### TLR4-mediated activation of dendritic cells *in vivo* requires INTS11

Since histological abnormalities were observed in spleen after INTS11 depletion (Fig. 6D), we performed flow cytometry to assess specific changes in spleenocytes. We found a significantly reduction in erythroid lineage in the spleenocytes isolated from *Ints11*^Δ*/*Δ^ mice (Fig. S7A), while myeloid and lymphoid lineages were affected to a small degree following *Ints11* deletion.

Dendritic cells (DC) in spleen are tasked with presenting antigens in the mammalian immune system (*30*). The mild reduction in myeloid lineage caused by *Ints11* deletion in spleen, provided an opportunity to investigate the responsiveness of myeloid-derived DCs to an inflammatory stimulus. Since DCs are activated in the response to foreign antigens such as lipopolysaccharide (LPS), a TLR4-mediated signaling stimulus, they presented a good model to assess Integrator’s role in activation of DCs gene expression programs(*31, 32*) (Fig. 7A).

**Figure 7.**
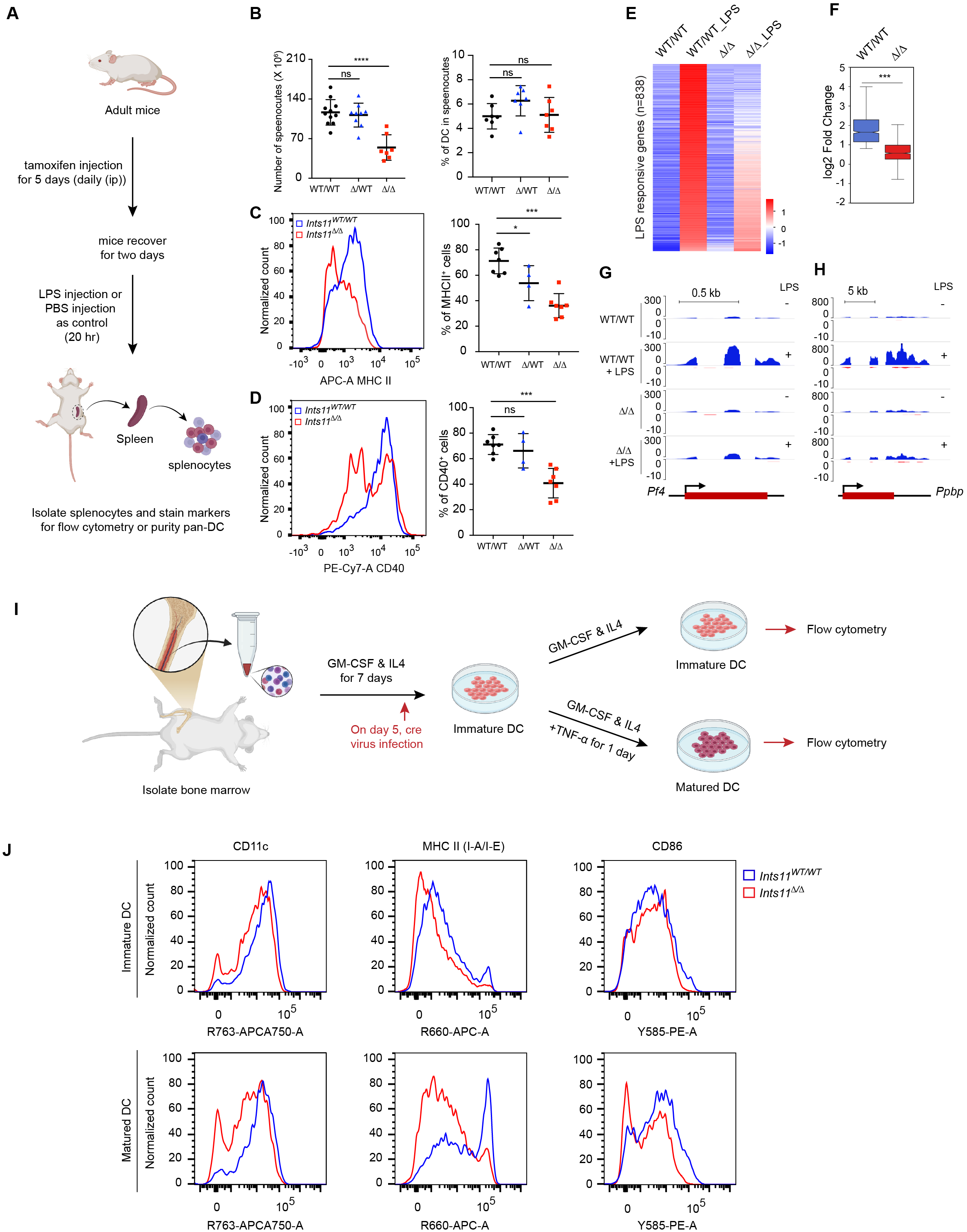
INTS11 is required for dendritic cell activation and maturation. **(A)** Experimental scheme for analysis of splenic dendritic cells. Adult mice were treated with TAM by intraperitoneal injection once daily for five consecutive days, allowed to recover for 2 days, and then injected with LPS or PBS as a control for 20 h. Spleens were collected, and splenocytes were analyzed by flow cytometry or used for purification of pan-dendritic cells (pan-DCs). **(B)** Quantification of total splenocyte numbers (left) and the percentage of dendritic cells among splenocytes (right) from *Ints11*^WT/WT^, *Ints11*^Δ/WT^, and *Ints11*^Δ*/*Δ^ mice. **(C)** Representative flow-cytometry histograms of MHC class II expression in splenic dendritic cells from *Ints11^WT/WT^* and *Ints11*^Δ/Δ^ mice following LPS stimulation (left), and quantification of MHC class II–positive dendritic cells across the indicated genotypes (right). **(D)** Representative flow-cytometry histograms of CD40 expression in splenic dendritic cells from *Ints11^WT/WT^* and *Ints11*^Δ/Δ^ mice following LPS stimulation (left), and quantification of CD40-positive dendritic cells across the indicated genotypes (right). **(E)** Heat map showing expression of 838 LPS-responsive genes in pan-DCs isolated from *Ints11^WT/WT^* and *Ints11*^Δ/Δ^ mouse spleens before and after LPS stimulation. Genes are ordered according to their transcriptional response to LPS. **(F)** Distribution of LPS-induced log2 fold changes across the 838 LPS-responsive genes in pan-DCs from *Ints11^WT/WT^* and *Ints11*^Δ/Δ^ mice. **(G and H)** Representative RNA-seq genome-browser tracks at the LPS-responsive *Pf4* **(G)** and *Ppbp* **(H)** loci in pan-DCs from *Ints11^WT/WT^* and *Ints11*^Δ/Δ^ mice before and after LPS stimulation. **(I)** Experimental scheme for generation and maturation of mouse bone marrow-derived dendritic cells (mBMDCs). Bone marrow cells were cultured with GM-CSF and IL-4 for 7 days, with Cre virus introduced on day 5. Cells were analyzed as immature BMDCs or further stimulated with TNF-α for 1 day to induce maturation before flow-cytometric analysis. **(J)** Representative flow-cytometry histograms showing CD11c, MHC class II (I-A/I-E), and CD86 expression in immature and TNF-α–matured BMDCs derived from *Ints11^WT/WT^* and *Ints11*^Δ/Δ^ mice.

**Figure S7.**
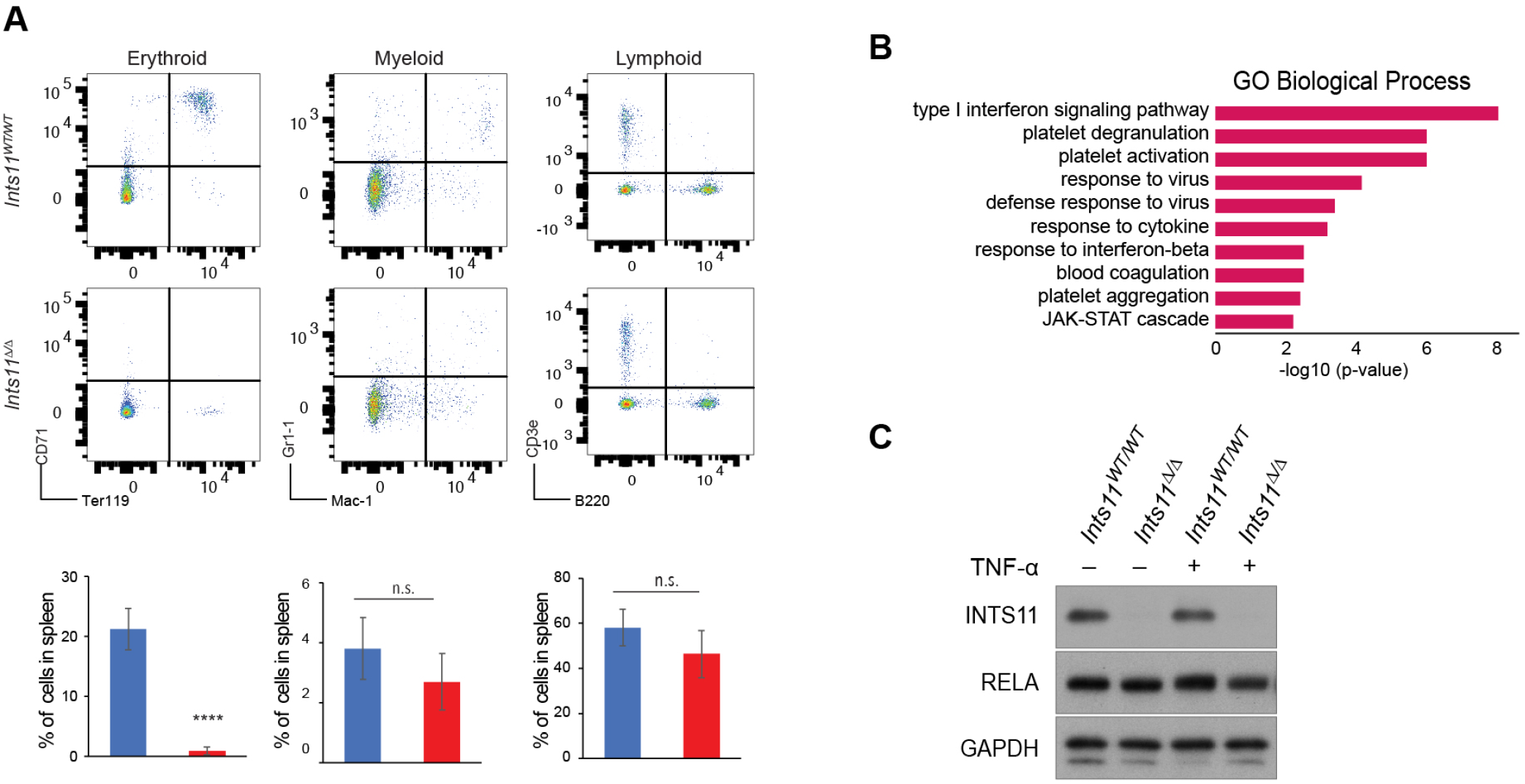
Additional characterization of INTS11-dependent dendritic cell activation and maturation. **(A)** Representative flow-cytometry plots (top) and corresponding quantification (bottom) of erythroid, myeloid, and lymphoid populations in spleens from *Ints11^WT/WT^* and *Ints11*^Δ/Δ^ mice. Erythroid, myeloid, and lymphoid populations were identified using CD71/Ter119, Gr-1/Mac-1, and CD3ε/B220, respectively. **(B)** Gene Ontology (GO) Biological Process enrichment analysis of genes up-regulated following LPS stimulation in dendritic cells isolated from adult mouse spleen. Enrichment significance is expressed as −log10(*P* value). **(C)** Immunoblot analysis of INTS11 and RELA in mBMDCs derived from *Ints11^WT/WT^* and *Ints11*^Δ/Δ^ mice before and after TNF-α stimulation. GAPDH served as a loading control.

We depleted INTS11 by treating mice with TAM and then challenged the animals with LPS to induce an immune response (Fig. 7A). While loss of INTS11 resulted in decreased number of splenocytes following homozygous *Ints11* deletion, it did not result in a change in the percentage of dendritic cells within the spleen (Fig. 7B). In contrast to homozygous deletion of *Ints11*, heterozygous mice did not display a significant decrease in the total number of splenocytes or percent of dendritic cells (Fig. 7B). While dendritic cells with wild type INTS11 express MHCII and CD40 as surface markers upon exposure to LPS, *Ints11* deletion resulted in a significantly diminished response (Fig. 7C-D). Indeed, analyses of pan-DC enriched cells using RNA sequencing revealed a profound loss of transcriptional signature associated with LPS-induced DC activation following *Ints11* deletion (Fig. 7E-H, 66% of differentially expressed LPS-induced genes were significantly attenuated; Fig. S7B, Table 5) These results highlight a critical role for INTS11 in activation of innate immune transcriptional response by the bacterial toxin LPS *in vivo*.

### Integrator is required for maturation of dendritic cells *in vitro*

A complementary *in vitro* model to activate DCs into a mature state is the use of TNF-α **to induce DC maturation from** mouse bone marrow-derived cells (DC)(*33, 34*). Therefore, we examined TNF-α-induced maturation of DC from wild type or homozygous *Ints11* deleted mouse bone marrow cells (Fig. 7I, Fig. S7C). Following TNF-α stimulation, wild type immature DC display enhanced MHCII and increased levels of CD86 and CD11c upon maturation (Fig. 7J). Deletion of *Ints11* abrogates MHCII expression, while the levels of CD86, and CD11c are concomitantly decreased (Fig. 7J). Collectively, our results pinpoint a role for Integrator in maturation of DC expressing antigen-stimulated gene expression signature in response to LPS *in vivo* and TNF-α stimulation in an *in vitro* model.

## Discussion

Stimulus-responsive transcription requires not only the arrival of transcription factors at chromatin but also the conversion of transient DNA encounters into stable, transcriptionally productive complexes. Our findings identify the Integrator complex as a critical determinant of this transition for NF-κB. We previously showed that Integrator is recruited to immediate-early genes after growth factor stimulation and coordinates transcription downstream of mitogen-activated protein kinase signaling (*23, 24*). Here, we extend this function to inflammatory signaling through the canonical NF-κB pathway. Together, these findings position Integrator as a broadly deployed regulator of rapid, stimulus-dependent transcription rather than a dedicated effector of a single signaling pathway.

Chromatin accessibility is an important determinant of NF-κB recruitment. Although a subset of NF-κB binding sites is embedded within nucleosomes and requires lineage-determining transcription factors to establish accessibility, most NF-κB sites reside within pre-accessible promoter and enhancer regions (4, 13, 14). Our results reveal an additional regulatory step operating after accessibility has been established. Integrator localized near NF-κB binding sites and was required for efficient NF-κB chromatin association after TNFα or LPS stimulation. Acute degradation of INTS11 was sufficient to disrupt NF-κB recruitment and transcriptional activation, arguing against an indirect consequence of prolonged Integrator loss. These findings define a post-accessibility checkpoint at which Integrator determines whether NF-κB engagement becomes sufficiently stable to generate a transcriptional response.

Single molecule tracking and fluorescence recovery after photobleaching provide a mechanistic basis for this function. Integrator markedly prolonged the chromatin residence of RELA molecules occupying a low-mobility, chromatin-associated state. Thus, Integrator functions as a chromatin-retention factor that converts short-lived NF-κB encounters into sustained binding events capable of supporting robust transcription. This interpretation is consistent with the loss of NF-κB occupancy detected by chromatin immunoprecipitation after Integrator depletion. Importantly, residence time is not simply equivalent to chromatin accessibility or steady-state occupancy; it represents a kinetic property of transcription-factor binding that can determine whether an enhancer recruits the machinery necessary for productive transcription. Our findings therefore establish NF-κB dwell time as a regulated step in inflammatory gene activation.

A central and unexpected finding is that this activity is independent of the canonical endonuclease function of Integrator. Catalytically inactive INTS11-E203Q fully restored NF-κB-dependent transcription, demonstrating that RNA cleavage is dispensable for this response. Integrator therefore exerts at least two mechanistically separable functions in transcription: a catalytic activity that regulates RNA polymerase II pause release and a structural activity that stabilizes transcription-factor association with chromatin. Whether Integrator anchors RELA through direct physical contact or stabilizes a larger enhancer-bound assembly remains to be resolved. Nevertheless, the association of Integrator with NF-κB, their genomic co-occupancy, and the rapid loss of RELA binding after INTS11 degradation support a model in which Integrator acts locally to stabilize productive NF-κB–chromatin interactions.

The physiological consequences of this mechanism extend across inflammatory stimuli, cell types, and species. Conditional loss of *Ints11* in adult mice profoundly impaired TLR4-dependent dendritic-cell activation, whereas Integrator deficiency attenuated LPS-induced transcription in mouse embryonic fibroblasts. Depletion of Integrator subunits in human cancer cells similarly suppressed TNFα-induced inflammatory gene expression. Integrator is therefore not merely associated with NF-κB target genes but is required for mammalian cells to translate inflammatory signals into appropriate transcriptional and cellular responses.

Collectively, our findings support a model in which Integrator coordinates two successive stages of inducible transcription: it first stabilizes signal-activated transcription factors at their regulatory elements and then promotes productive RNA polymerase II transcription. By coupling transcription-factor residence to polymerase activation, Integrator bridges enhancer recognition and transcriptional output. More broadly, these results suggest that the evolutionary emergence of Integrator in multicellular organisms provided a mechanism for converting transient extracellular signals into rapid and sustained gene-expression programs. Integrator thus represents a previously unrecognized regulatory layer in innate immunity, one that determines not simply whether NF-κB reaches chromatin, but whether it remains bound long enough to activate an inflammatory response.

## Acknowledgements

We thank the Shiekhattar lab for constructive discussions and suggestions for experimental design. We thank the Oncogenomics and flow cytometry shared resources at Sylvester Comprehensive Cancer Center for performing high-throughput sequencing and FACS analyses, respectively. This work was supported by funding from Sylvester Comprehensive Cancer Center P30CA240139 and grants R01 GM078455, and DP1 CA228041 from the National Institute of Health to R.S. Research reported in this publication was supported by the National Cancer Institute of the National Institutes of Health under Award Number P30CA240139. The content is solely the responsibility of the authors and does not necessarily represent the official views of the National Institutes of Health. S.S. is supported by the Leukemia Research Foundation, AFAR, the Japan Agency for Medical Research and Development, Merck, and the NIH grants 1S10OD030286-01 and P30CA01333047. B.A.G. is supported by the NIH grants R01AI118891 and P01CA196539.

## Author Contributions

R.S. and P.R.C. conceived and designed the overall project. P.K.R.C. performed RNA-seq, ChIP-seq, PRO-seq, RT-qPCR, ChIP-qPCR, SMT, FRAP, Immunostaining experiments. J.Y., O.L. and F Liu performed the mBMDC and MEFs, *in vivo* and *in vitro* experiments. F.B., H.G.D.S. and P.R.C. performed bioinformatics analyses. G.G. acquired and analyzed the STORM images. P.R.C., S.S., B.G. and F Lai performed ChIP-MS. R.S., P.R.C. and S.D. N. wrote the manuscript with critical feedback from all co-authors.

## MATERIALS AND METHODS

### Cell Lines

HeLa cells were cultured in Dulbecco’s Modified Eagle’s Medium (DMEM) supplemented with 10% FBS at 37°C with 5% CO2. Various HeLa cell lines, including HeLa-shCTRL, HeLa-shINTS4, HeLa-shINTS6, HeLa-shINTS9, HeLa-shINTS11 inducible KD clones, as well as shRNA-resistant N-terminal Flag-tagged wild-type (WT) or E203Q mutant INTS11 cell lines, were established as previously described(*20, 21*). The dTAG-INTS11 HeLa cell line used for acute depletion of INTS11 was generated as previously described(*35*). To assess their responsiveness to TNF-α or LPS, cells were treated with 10 ng/ml TNF-α or 200 ng/ml LPS for 1 hour.

### Antibodies and Reagents

RELA (Cell Signaling Technology-8242S), p50 (Cell Signaling Technology-3035S), INTS11(Bethyl Laboratories-A301-274A), INTS11 (Atlas Antibodies # HPA029025), INTS9 (Cell Signaling Technology-13945S), INTS4 (Bethyl Laboratories-92105), INTS6 (N-90,N-terminal, homemade), H1^0^ (Santa Cruz-sc-56694), RPB1-NTD (Cell Signaling Technology-14958), GAPDH (Cell Signaling Technology-5474S), BV421 Rat Anti-Mouse CD40 (BD Biosciences-562846), APC-Cy7 Hamster Anti-Mouse CD11c (BD Biosciences-561241), MHC class II (I-A/I-E) APC (R&D Systems-FAB6118A), and B7-2/CD86 PE (R&D Systems-FAB741P), TNFa (Sigma-T6674), LPS (Sigma-L2630), RNase A/T1 (Thermo Scientific-EN0551), DNase I (Promega-M6101), Turbo DNase (Invitrogen-AM1907), Biotin-11-NTPs (Perkin Elmer-NEL54(2/3/4/5)001), Dynabeads M-280 streptavidin (Invitrogen-11205), NEBNext Ultra II DNA library prep kit (NEBCat-E7645S), Truseq Stranded Total RNA library prep kit (Illumina-20020596).

MEFs were isolated from embryos of integrator conditional knockout mice. To induce integrator deletion, MEF cells were infected with retrovirus expressing Cre recombinase. Two days after infection, the cells were selected with puromycin for 3 days. To check the responsiveness to LPS, the cells were treated with 200ng/ml LPS for 1h.

### Tamoxifen-induced whole-body deletion of *Ints11*

To induce whole-body deletion of *Ints11* in adult mice, *Ints11*^flox/flox^ mice were crossed with Rosa26-CreERT2 mice to generate *Ints11*^flox/flox^;Rosa26-CreERT2^+/−^ animals. Adult mice aged 2–4 months received tamoxifen by intraperitoneal injection at 100 mg/kg once daily for five consecutive days. Cre-positive mice lacking the homozygous floxed *Ints11* genotype received the same tamoxifen regimen and served as controls. Animals were monitored daily for body weight, general health, and survival. To evaluate the effect of Ints11 deletion, mice received 5 doses tamoxifen were euthanized 7 days after initiation of tamoxifen treatment. Bone marrow, spleen, lung, liver, kidney, and brain tissues were collected for subsequent analyses. Deletion of *Ints11* was confirmed by RT-qPCR, which demonstrated reduced *Ints11* mRNA expression across the examined tissues. Tissue morphology was evaluated by hematoxylin and eosin staining, and hematopoietic and immune-cell populations in the bone marrow and spleen were characterized by flow cytometry.

### Analysis of dendritic cells derived from mouse bone marrow: *in vivo* and *in vitro* approaches

All animal works performed in this study were approved by the Institutional Animal Care and Use Committee at the University of Miami and conducted following institutional and national regulatory standards. For our *in vivo* study on LPS-induced dendritic cell maturation, we generated an inducible whole body knockout model by crossing our animal model with Rosa26-CreERT2 (Cre recombinase-estrogen receptor T2) mice (Jackson Laboratory, Stock No: 008463). Both control and Ints11 floxed mice were intraperitoneally injected with tamoxifen (75mg/Kg per day) over a period of five consecutive days to induce the deletion of Ints11. After a two-day resting period, the mice received an intraperitoneal injection of LPS (0.1mg/Kg, single dose). Twenty hours following the LPS injection, the animals were sacrificed, and spleens were collected. Spleenocytes were then isolated from the spleens. These isolated cells underwent two different processes: some were stained with surface markers for subsequent flow cytometry analysis, while others were subjected to a mouse pan-dendritic cell enrichment kit II (Stemcell Technologies, Cat #19863) to purify pan-dendritic cells for RNA-seq analysis.

For our in vitro mouse dendritic cell differentiation and maturation assay, we initiated the process by isolating bone marrow cells from both control and Ints11 floxed mice. These cells were then cultured in a differentiation medium supplemented with GM-CSF and IL-4, utilizing the Mouse Dendritic Cell Differentiation Kit provided by R&D Systems (Catalog Number: CDK008). This culture was maintained for a duration of five days to promote the differentiation of the cells into immature dendritic cells. Subsequently, we introduced a Cre recombinase-expressing retrovirus carrying a GFP reporter into the cell culture. On the seventh day of the experiment, TNF-α was introduced into the differentiation medium for a 24-hour period to induce dendritic cell maturation. Following this maturation step, the cells were subjected to staining to label their surface characteristics, after which they underwent flow cytometry analysis for further examination.

### Isolation of bone marrow and generation of derived macrophages

Mouse bone marrow–derived macrophages (mBMDM) were prepared from both control and Ints11 inducible knockout mice. In summary, bone marrow was flushed with PBS and cultured in differentiation medium enriched with 15% L929 cell-conditioned medium (LCM), which was added to complete DMEM. Cells were treated with 4-Hydroxytamoxifen or a vehicle for 5 days, starting on day 8, to induce Ints11 knockout. Subsequently, they were cultured without tamoxifen for an additional 2 days. On day 14, the differentiated cells were exposed to LPS for 60 minutes and then harvested for subsequent analysis.

### RNA-Seq and data analysis

Total RNA was extracted using Trizol reagent (Thermo Fisher Scientific, #15596026) following the manufacturer’s instructions. To eliminate genomic DNA, Turbo DNase treatment (Invitrogen, #AM1907) was applied. Total RNA-seq libraries were prepared using the TruSeq Stranded Total RNA library prep kit (Illumina, #20020596) with 500 ng of DNase-treated Input RNA. Sequencing was conducted on a NovaSeq 6000 or NextSeq 500 (Illumina) platform, provided by the Onco genomics Core Facility at the Sylvester Comprehensive Cancer Center, University of Miami Miller School of Medicine. All genome-wide experiments were performed as two independent biological replicates.

Raw fastq RNA-seq data were processed with Trimmomatic v0.32(*36*) and aligned to human genome (hg19 version) or mouse genome (mm10 version) using STAR aligner v2.5.3a(*37*) with default parameters and RSEM v1.2.31(*38*) to obtain expected gene counts against the human Ensembl (release 87) or mouse Ensembl (release 96).

Differential expression was determined in human (HeLa cells) for INTS11 shRNA (+Dox + TNF-α vs +Dox), GFP shRNA (+Dox + TNF-α vs +Dox) and siRNA against RELA (siRELA + TNF-α vs siRELA) and in mouse embryonic fibroblast (MEF) MEF WT (WT vs WT LPS) and MEF INTS11 KO (KO vs KO LPS) using DESeq2(*39*) and R v3.2.3. The genes with differential expression (TNF-α or LPS -responsive genes) were considered significant when the q-value was <0.05, fold change was >1.75, and TPM (Transcripts Per Million) was >1. For visualization on the UCSC Genome Browser, all tracks were CPM (count per million) normalized against the total number of usable reads in that data set using deepTools2(*40*).

### ChIP-seq, reChIP-seq and data analysis

ChIP was conducted in accordance with previously described(*41*) with slight modifications. For the INTS11 ChIP experiments, as depicted in Figure 2 and Figure S2 utilizing the Atlas INTS11 antibody, and the RNAPII ChIP, cells were initially cross-linked with 1% formaldehyde for 10 minutes at room temperature. The cross-linking reaction was quenched by incubating the cells with 125 mM glycine for 5 minutes before harvesting. In the INTS11 ChIP and reChIP experiments illustrated in Figure 5 and Figure S5, utilizing the Bethyl INTS11 antibody and involving RELA ChIP, samples underwent initial cross-linking with Cross-link Gold (Diagenode C01019027) following the manufacturer’s instructions before proceeding to the formaldehyde fixation step. The pellet was resuspended in ChIP lysis buffer (20 mM Tris-HCl, 50 mM NaCl, 0.1% SDS, 0.5% Triton X-100, 1 mM EDTA) and sonicated in a Covaris M220 instrument under the following conditions: duty cycle 10, peak incidence power 140, cycles per burst 200, until chromatin fragments of 150-400 bp were achieved. After sonication, chromatin was clarified by centrifugation at 16,100 ×g for 15 minutes at 4°C, and the supernatant was transferred to a tube and adjusted to a final concentration of 150 mM NaCl. Protein A/G magnetic beads bound with antibodies were incubated with sonicated chromatin overnight. The following day, beads were subjected to two washes with each of the following buffers: Mixed Micelle Buffer (20 mM Tris-HCl, 150 mM NaCl, 1% Triton X-100, 0.2% SDS, 5 mM EDTA, 65% sucrose), Buffer 500 (25 mM HEPES, 500 mM NaCl, 1% Triton X-100, 0.1% Na deoxycholate, 10 mM Tris-HCl, 1 mM EDTA), LiCl/detergent wash (10 mM Tris-HCl, 250 mM LiCl, 0.5% Na deoxycholate, 0.5% NP-40, 1 mM EDTA), and a final wash was performed with TE buffer. Finally, beads were resuspended in elution buffer (10 mM Tris-HCl, 1 mM EDTA, 1% SDS) and incubated at 65°C for 20 minutes; elution was repeated twice. For reChIP-seq analysis, the ChIP-eluted material was subsequently employed for reChIP experiments. The eluted material was adjusted to a final concentration of reChIP buffer, consisting of 20 mM Tris-HCl, 150 mM NaCl, 0.1% SDS, 0.5% Triton X-100, and 1 mM EDTA. Protein A/G magnetic beads prebound with specific antibodies were incubated with this eluted material. The washing and elution steps were performed following a procedure similar to that used in standard ChIP-Seq experiments. The samples were subjected to overnight incubation at 65°C to reverse the cross-linking, along with an untreated input sample, which represented 10% of the starting material. The following day, eluates were treated with RNase A and incubated at 37°C for 1 hour, followed by Proteinase K incubation for 2 hours at 55°C. Subsequently, a phenol:chloroform:isoamyl alcohol extraction was performed, followed by ethanol precipitation. The resulting DNA was resuspended in 50 μL TE buffer and utilized for either qPCR or sequencing analyses. ChIP-qPCR assays were carried out with region-specific primers, and data were collected using a CFX96 real-time system (BioRAD) with iTaq universal SYBR green supermix. ChIP-seq and reChIP-seq libraries were constructed using the NEBNext Ultra II DNA library prep kit (New England Biolabs, #E7645S) and subsequently sequenced on a NovaSeq 6000 or NextSeq 500 (Illumina) platforms. All genome-wide experiments were performed as two independent biological replicates.

Raw FASTQ data were processed with Trimmomatic v0.32 to remove low-quality reads and then aligned to the human genome hg19 or mouse genome mm10 using STAR aligner v2.5.3a(*37*). We used deepTool2(*40*) to generate normalized bigwig. Peaks were called using MACS2.1.2 with the following parameters --shift 75 --nomodel --extsize 200 -q 0.05 for narrow peaks (RELA). Whole-cell extract input from the corresponding cell lines were used as controls. The R Bioconductor package DiffBind(*42*) was used for identifying sites that are differentially bound between sample groups. In human the groups are: HeLa cells RELA Control vs RELA Control TNF-α and RELA WT Dox TNF-α vs RELA EQ Dox TNF and for mouse macrophages WT CTRL vs WT LPS and WT LPS vs EQ LPS. The differential binding analysis was performed with DESeq2. In Hela cells (RELA Control vs RELA Control TNF-α), we used the following cutoff fold change > 2 and q < 0.05 additional to and overlap top INTS11 + TNF-α peaks (n=7521) and in mouse macrophages (WT CTRL vs WT LPS), the cutoffs were fold change > 4 and q < 0.0001 (n=12090). We open a window of 20 kb (-/+ 10 kb) from the TSS of the 201 TNF-α responsive genes detected by RNAseq, and used bedtools intersect function to identify 406 RELA peaks in those genes. We used deepTool2(*40*) to generate all heatmaps.

### Rapid PRO-seq (rPRO-seq) and data analysis

rPRO-seq was performed as previously described(*35, 43*). 10×10^6 HeLa nuclei were mixed with 0.5 x10^6 spike-in Drosophila S2 cell nuclei. Nuclear run-on assays were then performed with 25 μM Biotin-11-ATP/GTP/CTP /UTP (PerkinElmer) for 3 min at 30 °C and RNA was fragmented. Biotinylated-RNA was separated with M280 Streptavidin Dynabeads (Thermo Fisher). Following adaptor ligation, reverse transcription, and PCR amplification. DNA Libraries were size selected by AMpure XP (Beckman Coulter) and sequenced by NovaSeq 6000 system (Illumina) with single-read runs. Raw fastq data were processed as described previously (*21*). Briefly rPRO-seq reads were trimmed by Cutadapt 1.14(*44*) and Trimmomatic v0.32(*36*) and aligned to the human genome (hg19) genome or the drosophila genome (dm3) by bowtie 1.1.2(*45*) rPRO-seq signal was normalized by the number of reads mapped to spike-in dm3 genome, and then converted to bigwig data, which were used for downstream analyses. Quantification of nascent transcription per region was generated with an in-house script based on pyBigWig http://dx.doi.org/10.5281/zenodo.45238) used to extract the normalized reads (CPM) per position from the bigwig files with a bin size of 1 bp upon merging the signal across two replicates. Read density was calculated per gene (longest isoform) and the distributions represented as boxplots. Two sample Kolmogorov-Smirnov nonparametric test (KS-test) was applied to determine if two datasets (for instance shRNA INTS11 vs. shGFP upon TNF induction) differ significantly by comparing their cumulative distributions (p-value <= 0.05).

### Stochastic optical reconstruction microscopy (STORM)

Cells were seeded on ibidi micro-slide 4 wells, fixed in 4% formaldehyde for 10 min at room temperature. Cells were then permeabilized in 0.5% Triton-X-100 for 15min, blocked with Normal goat serum overnight, and incubated overnight at −20 C with the primary antibody (H3) at 1:500 dilution and RELA (1:300 dilution). Finally, secondary antibodies (Novus Biologicals-NBP1-75398JF646 anti-rat and Alexa Fluor 568 anti-rabbit (Invitrogen - A10551) were applied to the samples at a dilution of 1:500 for 30 min at room temperature.

Imaging experiments were done with a Nikon eclipse Ti2 microscope equipped with Nikon Instruments(N-STORM). Two color dSTORM imaging was labeled with both Janelia 646, and Alexa 568 secondary antibodies. MEA STORM imaging buffer was applied to samples prior to imaging. Images were acquired at 512 x 512 pixels, at a frequency of 20ms for a total of 5000 frames per filter. Images were acquired using a 100x, 1.49 NA oil immersion objective, and imaged using a Hamamatsu C11440 ORCA-flash 4.0 camera. STORM localization analysis was carried out with ImageJ, thunderstorm plugin (1.3-2014-11-08). Data was fitted with a Gaussian PSF model using weighted least squares estimation. Molecule files were then exported from Image J to be further analyzed using Coloc-Tessler(*46*). A two-tailed t test was performed using GraphPad Prism 5 to determine significance of colocalization.

### Co-immunoprecipitation (CoIP)

HeLa cells were treated with TNF-α for 60 minutes. Subsequently, cell extracts were prepared using lysis buffer (20 mM Tris-HCl, pH 7.9, 10 mM KCl, 1.5 mM MgCl2, 150 mM NaCl, 1% Triton X-100, and 0.5 mM phenylmethylsulfonyl fluoride [PMSF]). Antibodies (INTS11 or IgG) pre-bound to Protein A/G magnetic beads were incubated with cell extracts overnight at 4°C. To prevent potential indirect protein interactions mediated by nucleic acids, the cell extracts were treated with either RQ1 DNase I or RNase A/T1 during the immunoprecipitation process. The beads were then subjected to washes as follows: two washes with wash buffer (50 mM Tris-HCl, pH 7.9, 0.1% Triton X-100, and 0.5 mM PMSF) containing 500 mM NaCl, two washes with wash buffer containing 250 mM NaCl, and one wash with wash buffer containing 150 mM NaCl.

### Single-molecule imaging and analysis

Live-cell single-molecule tracking (SMT) of RELA-HaloTag was performed using a Nikon N-STORM system built on a Ti2 inverted microscope and operated in total internal reflection fluorescence (TIRF) mode. Images were acquired through an SR HP Apo TIRF 100×H oil-immersion objective (NA 1.49) using a Flash4.0 sCMOS camera with 1 × 1 binning. For residence-time measurements, movies consisted of 300 frames acquired at an approximately 100-ms frame interval with a 30-ms exposure time. Images were collected at 256 × 256 pixels with a calibrated pixel size of approximately 0.065 μm/pixel. Imaging settings were maintained consistently across the experimental conditions being compared. Single-molecule localization and trajectory reconstruction were performed using the MATLAB routine MatlabTrack_v5.03, following previously described SMT procedures (*47, 48*). Individual RELA-HaloTag molecules were localized in consecutive frames and linked into trajectories using the same analysis parameters across experimental conditions. RELA molecules were classified into slow-fraction, fast-fraction, and unbound populations. For residence-time analysis, RELA-HaloTag trajectories were analyzed following previously described procedures (*47, 48*), and residence times were determined for the slow- and fast-fraction populations. The same trajectory-selection and analysis parameters were applied across all experimental conditions.

### Fluorescence recovery after photo bleaching (FRAP)

FRAP experiments were performed on Hela cells transiently transfected with GFP and GFP-RELA 48 h before FRAP. For data acquisition, cells were analyzed with the Nikon eclipse Ti2 microscope equipped with Nikon Instruments (NSTORM) using a 100× oil immersion (NA 1.40) objective, at zoom 5, at 37^°^C. Selected areas were bleached using the 488 nm Argon laser set with maximum laser intensity and images were acquired every 250 ms for 181 cycles. The FRAP curve and half time of recovery (t***_1/2_***) were obtained using easyFRAP-web, according to the website’s instructions(*49*). Data representation was performed in GraphPad Prism 8.

### Immunostaining

Cells were fixed in 4% paraformaldehyde (PFA) in PBS for 20 min at room temperature and washed three times with PBS. Cells were permeabilized with 1% Triton X-100 in PBS for 20 min, washed with PBS, and blocked with 3% protease-free bovine serum albumin (BSA; Roche, #03117332001) for 1 h at room temperature. Primary antibodies were diluted 1:400 in blocking solution and incubated overnight at 4°C. Following three washes with PBS, cells were incubated with the appropriate secondary antibodies for 30 min at room temperature and washed with PBS. Nuclei were stained with Hoechst (8 μg/mL in PBS) for 5 min, followed by three PBS washes. Images were acquired using a Leica THUNDER Imager microscope.

### siRNA transfection

siRNA transfection was performed using Lipofectamine RNAiMax transfection reagent (Invitrogen, catalog no. 13778-100) according to the manufacturer’s instructions. The siRNAs were purchased from Ambion, negative control siRNA [ambion-4390843]), siRNAs against RELA [ambion-s11915]. All the PCR primer sequences are listed in the supplementary Table 6.

## Data Availability

All of the genome-wide data of this study have been deposited in the NCBI Gene Expression Omnibus (GEO) database. The accession number for the raw and processed data reported in this paper is GEO: GSE244770.

